# Frequency dependent growth of bacteria in living materials

**DOI:** 10.1101/2022.02.22.481564

**Authors:** Daniel D. Lewis, Ting Gong, Yuanwei Xu, Cheemeng Tan

**Author notes:** Equal contribution (Daniel D. Lewis, Ting Gong).

## Abstract

The fusion of living bacteria and man-made materials represents a new frontier in medical and biosynthetic technology. However, the principles of bacterial signal processing inside three dimensional and fluctuating environments of synthetic materials remain elusive. Here, we study bacterial growth in a three-dimensional hydrogel. We find that bacteria expressing an antibiotic resistance module can take advantage of ambient kinetic disturbances to improve growth while encapsulated. We show that these changes in bacterial growth are specific to disturbance frequency and hydrogel density. This remarkable specificity is consistent with stochastic resonance theory, which we leverage to explain how bacteria can integrate spatial and temporal information to control growth. This research provides a quantitative foundation for the control of living materials and a systematic framework towards understanding bacterial information processing in three-dimensional environments.

## Introduction

The integration of designer microbes with synthetic materials has birthed a new class of customizable, environmentally-responsive living materials (Balasubramanian et al., 2019; Cao et al., 2017; Gilbert et al., 2021; Gonzalez et al., 2020; Hay et al., 2018; Manjula-Basavanna et al., 2021). Genetic circuits inside bacteria can be designed to produce enzymatic or structural components based on intra- or extra-cellular cues. Through expression regulation mediated by genetic circuits, designer microbes can control self-assembly (Cao et al., 2017) and adaptation (Gilbert et al., 2021; Gonzalez et al., 2020; Hay et al., 2018) in synthetic materials. Designer microbes are often incorporated into hydrogels to form engineered living materials (Chyzy and Plonska-Brzezinska, 2020). Hydrogels’ physical properties, such as stiffness, density, and viscosity, play an important role in the metabolic activity and growth rate of encapsulated cells (Gilbert et al., 2021; Manjula-Basavanna et al., 2021). While the interactions between hydrogels and cellular growth can be exploited to control self-assembly (Cao et al., 2017), they can also impart metabolic heterogeneity. Metabolic heterogeneity of encapsulated cells in synthetic materials is known to impede their growth and genetic circuit activity (Alvarez et al., 2009; Pabst et al., 2016; Wessel et al., 2013). To expand the utility and range of biological functions in engineered living materials, there is a critical need to understand and control the growth of synthetically encapsulated microbes.

Foundational work on natural bacterial mechanotransduction has shown that cells are capable of integrating complex environmental inputs into decisions that reshape cellular physiology (Dufrene and Persat, 2020; Taute et al., 2015). However, these studies have also revealed gaps in our understanding of how bacteria integrate temporally and spatially heterogeneous signals into beneficial phenotypic changes (Kandemir et al., 2018; Sanfilippo et al., 2019). There have been explorations of the connections between spatial structure and information processing in microbial colonies (Alnahhas et al., 2019; Bittihn et al., 2020; Dal Co et al., 2020; Gupta et al., 2020; Kong et al., 2017; Kussell and Leibler, 2005; van Vliet et al., 2018; Zhang et al., 2021), but these studies have not addressed the information processing of microbes embedded in three-dimensional synthetic materials. Along this line, studies using engineered cells suggest that phenomena like Turing patterns may facilitate collective decision-making based on spatial and temporal information (Cao et al., 2016; Karig et al., 2018). Do unique information-processing principles govern microbial behavior in three-dimensional materials? If so, could those principles be used to control the growth of microbes encapsulated in synthetic materials?

Here, we study the capacity of bacteria to capitalize on rapid periodic disturbances in a synthetic hydrogel. Rapid periodic disturbances are common in complex 3D environments that are key targets for engineered living materials (Nguyen et al., 2018), such as cardiac tissue, airways, and intestines. We use an antibiotic resistance gene to explore how the metabolic trade-off of a synthetic genetic module affects bacterial growth in a spatially heterogeneous and temporally fluctuating environment. This work uncovers a counterintuitive phenomenon where the frequency of rapid periodic disturbances optimizes the growth of antibiotic-resistant bacteria encapsulated in hydrogel. A mathematical model recapitulates this frequency-dependent optimization of growth by proposing that cells can leverage stochastic resonance (McDonnell and Abbott, 2009; Paulsson et al., 2000; Zeng et al., 2009) to integrate spatial and temporal information to survive in synthetic materials.

## Results

### Bacteria exhibit fluctuating ATP metabolism in periodically perturbed hydrogel

Spatial heterogeneity caused by hydrogel encapsulation can disrupt cellular metabolism underlying signal propagation (Shao et al., 2017). However, it was unknown whether these disruptions in metabolism could prevent a cell from responding to rapid fluctuations in its environment. To address this unknown, we grew bacteria in a culture system where we could modulate the heterogeneity of the media, periodically perturb the cells, and measure the subsequent changes in intracellular signals (**Fig. 1A**). *Escherichia coli* DH5*α* was used as a basal strain, cultured in freshly-prepared LB (Luria Broth) media and hydrogel. Encapsulation conditions were controlled by dissolving hydrogel in media by autoclaving, then storing the media at 37°C until use (Methods-Section A).

**Figure 1:**
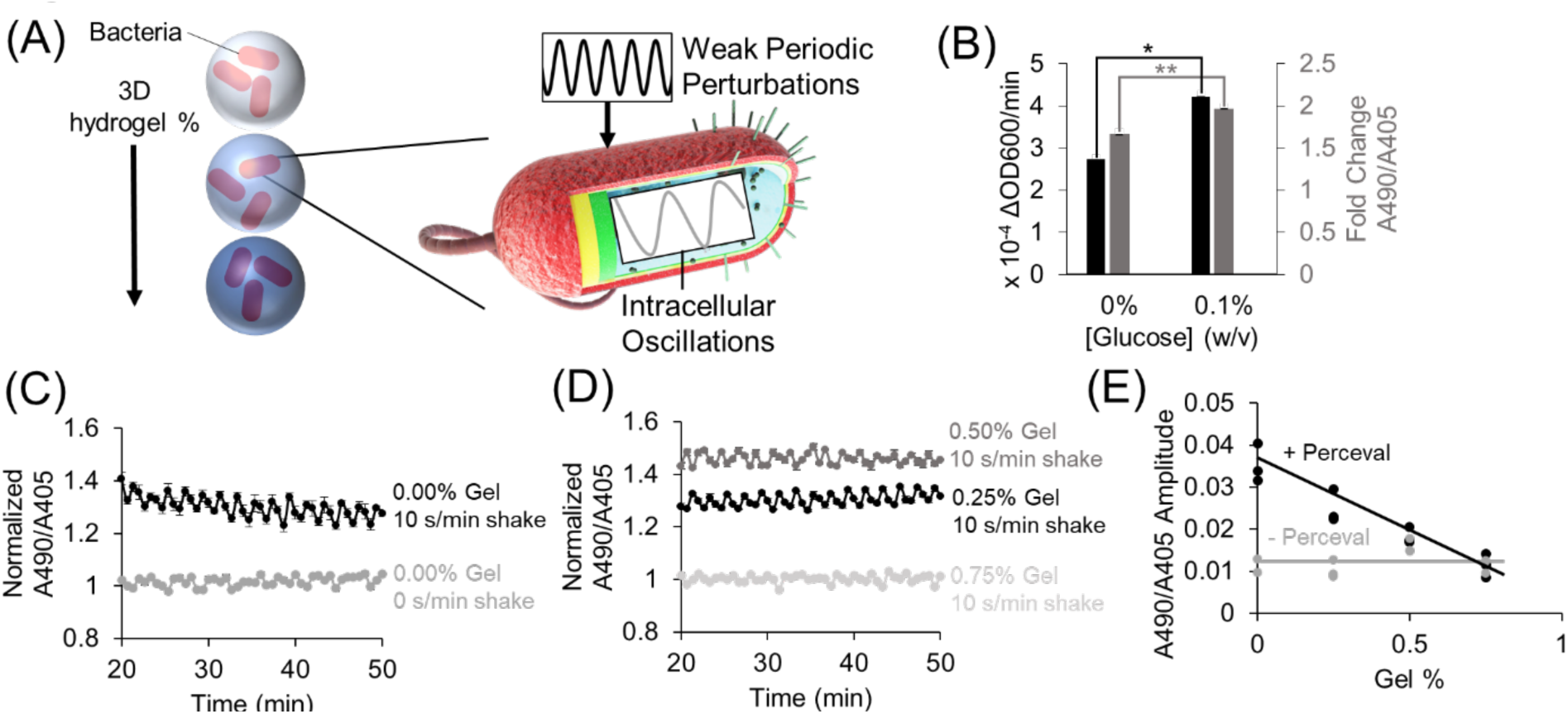
Periodic environment perturbations can propagate to intrabacterial metabolite oscillations despite hydrogel encapsulation. (A) Cartoon depicting bacteria encapsulated in varying densities of hydrogel being subjected to external kinetic perturbation. Experiment assays whether external perturbations propagate to internal fluctuations in cellular metabolites. (B) Bacterial growth rate (black bar) and their perceval expression (grey bar) in M9 minimal medium (0 and 0.1% glucose, respectively). Difference between bacterial growth rates is significant (*p*-value<0.05, indicated by asterisk). Difference between perceval values is significant (*p*-value<0.005, indicated by two asterisks). Bar height represents mean values, and error bars represent standard error of the mean. Mean and SEM are calculated using four biological replicates. (C) Intrabacterial ATP/ADP ratio over time with 0 or 10 s/min shaking. Absorbance at 490 nm to absorbance at 405 nm ratio represents ATP/ADP ratio. Basal fluctuation in A490/A405 without shaking is likely caused by the mechanical movement of the platereader to read the samples. Points represent mean values, and error bars represent standard error of the mean. Mean and SEM are calculated using four biological replicates. (D) Intrabacterial ATP/ADP ratio at different hydrogel densities (0.25%-0.75% gel). Absorbance at 490 nm to absorbance at 405 nm ratio suggests that periodic perturbation of hydrogel encapsulated cells can propagate to changes in intracellular metabolites. Points represent mean values, and error bars represent standard error of the mean. Mean and SEM are calculated using four biological replicates. (E) Amplitude of ATP/ADP fluctuations in *E. coli* DH5α cells (grey line) and Perceval cells (black line) at different gel concentrations (0-0.75%). Results show that increasing hydrogel concentration reduces amplitude of ATP/ADP ratio fluctuation. Points represent individual replicates, line highlights trend. At least two replicates per condition.

To monitor changes in intracellular metabolites, bacteria were grown expressing the metabolic reporter known as Perceval (Berg et al., 2009) (Methods-Section B&C). Perceval is designed to fluorescently report the ratio of ATP to ADP. Increased bacterial growth rate (**Fig. 1B**, black bar, *p*-value: 0.02) correlated with an increased ratio of absorbance at 490 nm to absorbance at 405 nm (**Fig. 1B**, grey bar, *p*-value: 0.001). Since the ratio of A490/A405 corresponds to the ATP/ADP ratio, these results suggest that the Perceval reporter can serve as a proxy for cellular bioactivity. These results are consistent with the previous use of ATP as a reporter of metabolic activity (Lopatkin et al., 2019).

Next, bacteria expressing Perceval were cultured in a periodically-shaken environment and changes in the ATP to ADP ratio were measured over time at different gel concentrations. In cultures shaken for 10 s/min, oscillations occurred in the A490/A405 ratio (**Fig. 1C** black). Some A490/A405 fluctuation was observed in the absence of shaking (**Fig. 1C** grey), but this is likely due to the mechanical movement required to measure different wells every two minutes. When hydrogel-encapsulated bacteria were shaken for 10 s/min, oscillation in the A490/A405 ratios was also recorded (**Fig. 1D** black, dark grey). These results challenge the assumption that heterogeneous conditions eliminate the dynamic responsiveness of intracellular metabolites to environmental conditions. High concentrations of gel were able to repress oscillations in the A490/A405 ratio (**Fig. 1D** light grey). These results suggest that sufficiently concentrated environmental hydrogel can dampen short-term cellular metabolic responses. A power spectral analysis (**Fig. 1E**, **Supp. Fig. 1** 2^nd^ Column) revealed that Perceval-expressing cells at 0.25% gel experienced significantly higher oscillation amplitude (*p*-value: 0.01) than background fluctuations in A490/A405 measurements. We observed similar periodicity in bacteria expressing a luminescent reporter under periodic and heterogeneous conditions (**Supp. Fig. 2**, Methods-Section D). These results demonstrate that rapidly oscillating physical perturbation of hydrogel-encapsulated bacteria can propagate to intracellular metabolic changes (**Fig. 1C-E**, **Supp Fig 1-2**).

### Bacterial gene expression and growth rate can be controlled by exogenous hydrogel

To confirm that the hydrogel used in this study imparts heterogeneity on growing cells, bacterial growth in a 3D hydrogel was measured. Bacterial density was recorded for two hours at a range of different hydrogel concentrations (**Supp Fig. 3A-D** top rows). Bacterial density at two hours was used to compare growth between cultures, serving as a surrogate reporter of cellular bioactivity (**Fig. 2A**). The two-hour time point was chosen because the rate of change in OD600 decreased after two hours. The results showed that increasing gel concentration from 0% to 0.75% (weight/volume) caused a 1.8 fold reduction in final bacterial density (**Fig. 2A**). This suggests that DH5*α* cells grew slower with increasingly dense hydrogel, resulting in reduced growth.

**Figure 2.**
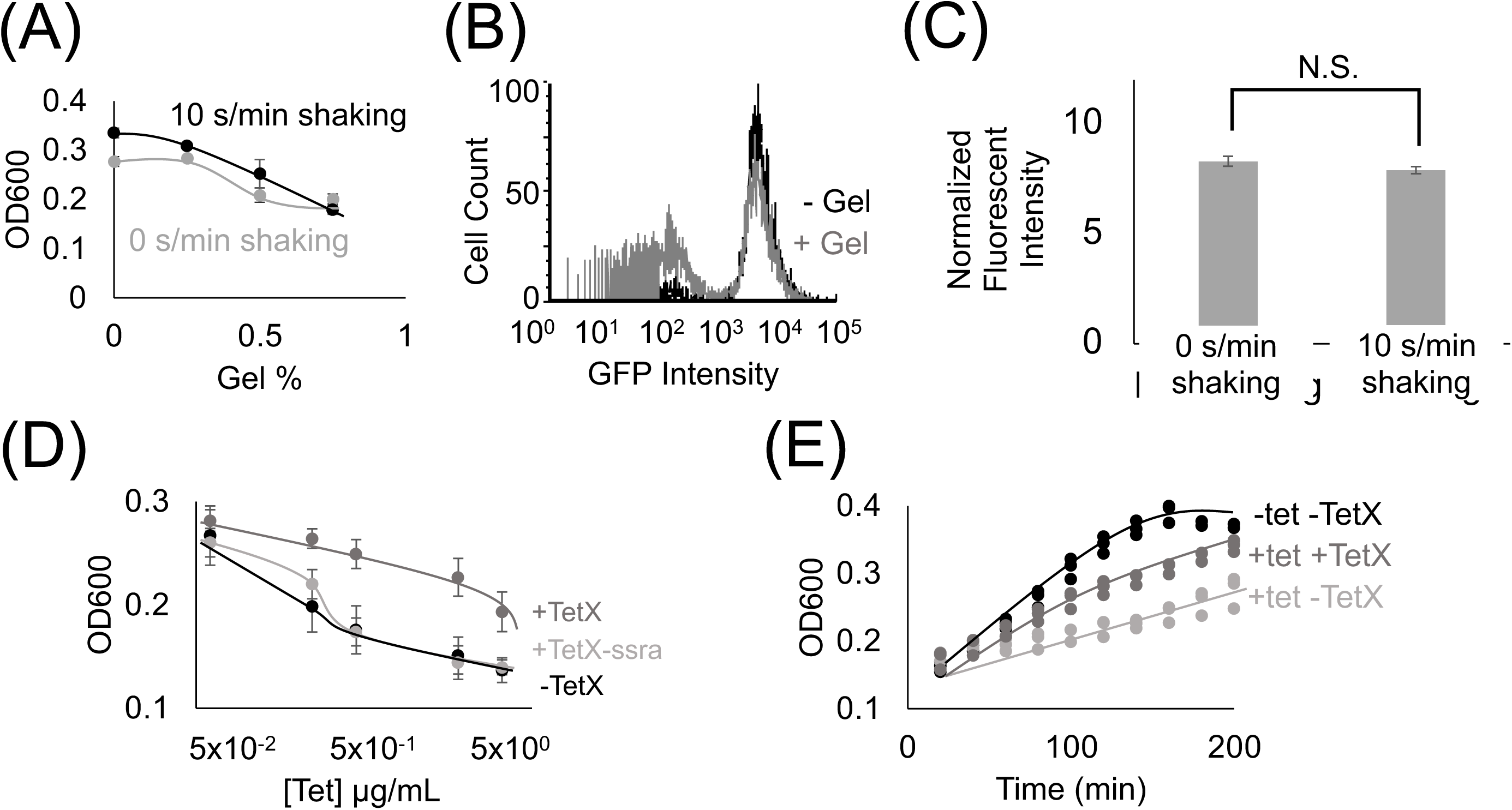
Hydrogel encapsulation imparts gene expression heterogeneity. (A) Cell density in different gel concentration environments at 2 hr. 0%, 0.25%, 0.5% and 0.75% gel density with 10 s/min shaking (black line) or with no shaking (grey line). Increased hydrogel concentration reduces bacterial growth. Points represent mean values, and error bars represent standard error of the mean. Mean and SEM are calculated using six biological replicates. (B) Fluorescence intensity distribution of cells expressing TetX-GFP fusion protein in media containing 0% (black line) and 0.5% low-melting-temperature agarose (grey line). Results demonstrate that hydrogel-encapsulation limits the expression of TetX-GFP in a subset of bacteria. One of two replicates shown for each condition, refer to Supp. Fig. 4 for both replicates. (C) Fluorescence intensity of TetX-GFP normalized by OD600 of cells after two hours of growth with either 0 s/min or 10 s/min shaking. Mean expression levels are not significantly different. Two-tailed, equal variance t-test returns *p*-value = 0.189. N.S. stands for ‘not significant’. Bar height represents mean values, and error bars represent standard error of the mean. Mean and SEM are calculated using eight biological replicates. (D) Cell density of resistant and non-resistant DH5*α* cells under different concentrations of tetracycline. +TetX-ssra (light grey line), +TetX (grey line) and -TetX (black line). Results show that TetX rescues bacterial growth from tetracycline repression. Points represent mean values, and error bars represent standard error of the mean. Mean and SEM are calculated using six biological replicates. (E) Cell density of resistant bacteria (TetX) in different tetracycline (tet) over time. 0.625 µg/mL tetracycline on DH5*α* cells without TetX (light grey line), 0.625 µg/mL tetracycline on DH5*α* cells without TetX (grey line) and 0 µg/mL tetracycline on DH5*α* cells without TetX (black line). Results show that tetracycline inhibition of cellular growth is reflected by decreased rates of growth over time, and rescue by TetX is reflected by increased rates of growth over time. Points represent individual replicates. Six replicates per condition at each time point.

To establish the effect of exogenous hydrogel on gene expression modules, DH5*α* cells constitutively expressing GFP (Methods-Section B) were grown in media with 0.5% gel (w/v) or 0% gel. Single-cell expression data from DH5*α* cells grown in the presence of hydrogel showed that a secondary population of low-fluorescence cells appeared (**Fig. 2B**, **Supp Fig. 4**, Methods-Section E). This bimodal expression distribution suggests that hydrogel encapsulation imparts heterogeneous growth conditions that stochastically limit gene module expression levels, splitting the cellular population into “on” and “off” states. Taken together, hydrogel-mediated reductions in growth and gene expression are consistent with previous work on microbes in heterogeneous environments (Alvarez et al., 2009; Pabst et al., 2016; Wessel et al., 2013). Notably, changes in periodic perturbation did not significantly alter population-level fluorescence measurements (**Fig. 2C**, **Supp Fig. 5**). This observation could be explained by periodic perturbation inducing offsetting changes in gene expression rate and protein dilution rate.

### Engineered genetic module imparts functionality on encapsulated bacteria

To test the effect of protein expression on bacterial growth during encapsulation, a genetic module that constitutively expresses TetX was constructed (Ghosh et al., 2015) (Methods-Section A). We validated the efficacy of this TetX module by growing resistant and non-resistant bacteria at a series of tetracycline concentrations, demonstrating that TetX expression allows cells to grow to a 1.5-fold greater final density in the presence of 5 μg/mL tetracycline (**Fig. 2D**). This change in density is also reflected over time (**Fig. 2E**). To confirm that the protein product of the module (i.e. TetX) was imparting antibiotic resistance, an unstable version of TetX was constructed by fusing a degradation tag to TetX (Gottesman et al., 1998). The unstable protein either imparted very weak tetracycline resistance or no detectable tetracycline resistance (**Fig. 2D**). These results establish that TetX expression, rather than nonspecific protection from the TetX module, allows cells to effectively resist tetracycline (**Fig. 2D-E**).

### Bacteria exhibit emergent growth in a 3D fluctuating hydrogel

To examine the population-level consequences of functional protein expression for bacteria inside a periodically-perturbed hydrogel, the growth of DH5*α* cells constitutively expressing TetX was measured over time (**Fig. 3A,** Methods-Section F). 0, 0.3125, or 0.625 µg/mL tetracycline was supplemented in the media. To quantify relative growth, the final density of cells expressing the TetX module was normalized by initial density, and the growth of naïve cells cultured under the same conditions. Under all growth conditions, larger concentrations of tetracycline increased the relative growth of TetX-expressing cells. The significance of increases or decreases in relative growth between cells at different concentrations of hydrogel were assayed using a Bonferroni multiple comparisons test (Methods-Section G). This statistical test was used to classify trends as increasing, biphasic, decreasing, or constant (**Supp Fig. 6**).

**Figure 3.**
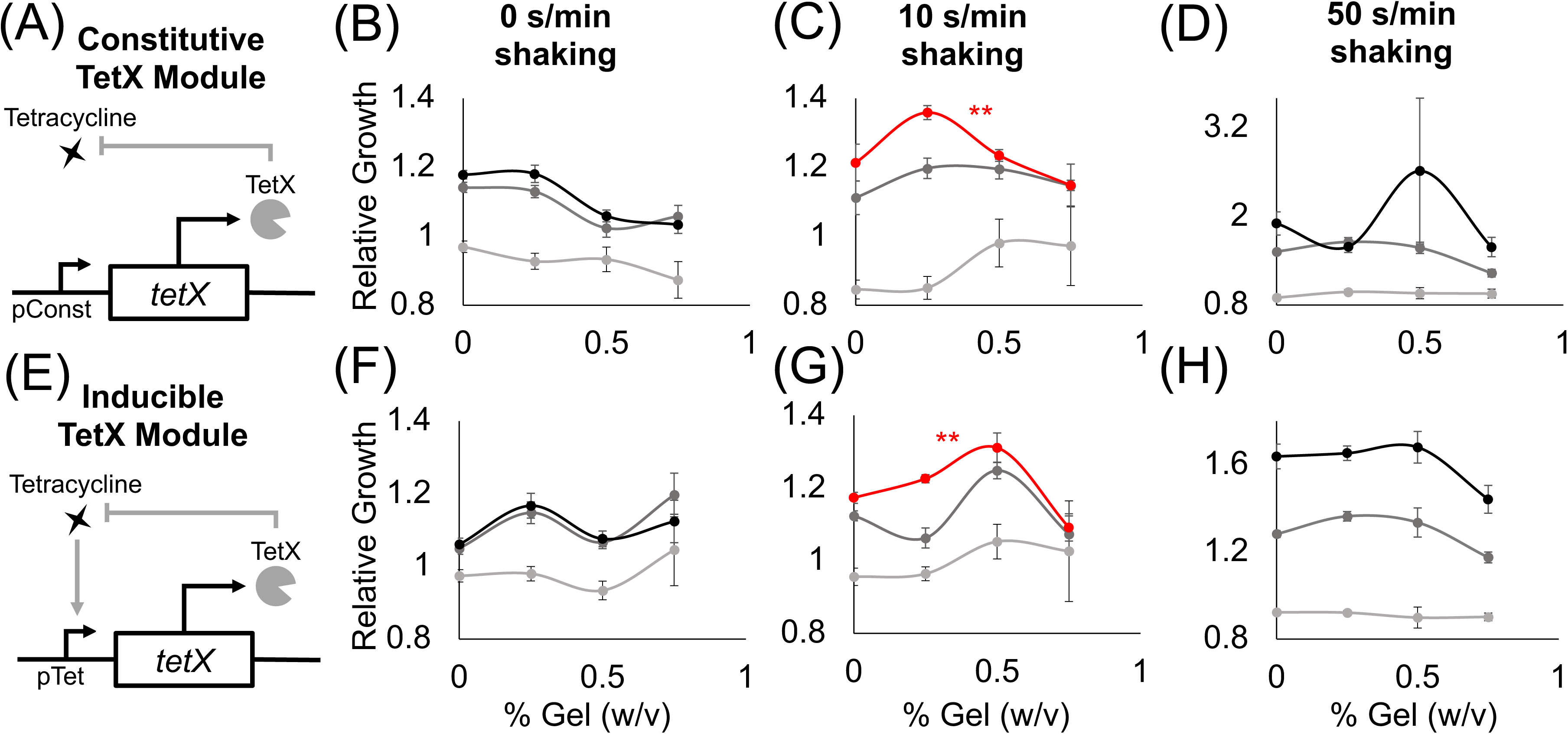
A minimal antibiotic-resistance module enables frequency-dependent response of bacterial growth in hydrogel. (A) Abstract depiction of tetracycline degradation by constitutive TetX expression module. (B-D) Relative growth of *E. coli* cells constitutively expressing TetX without shaking (B), with 10s shaking (C), with 50s shaking (D). Light grey is 0 µg/mL tetracycline, dark grey is 0.3125 µg/mL tetracycline, black is 0.625 µg/mL tetracycline. The red line represents a statistically significant biphasic curve as judged by a Bonferroni multiple comparisons test (treated with 0.625 µg/mL tetracycline). Points represent mean values, and error bars represent standard error of the mean. (B) and (C) Mean and SEM are calculated using six biological replicates. (D) Mean and SEM are calculated using four biological replicates. (E) Abstract depiction of feedback between bacterial growth and inducible TetX expression module when feeding tetracycline. (F-H) Relative growth of *E. coli* cells inducibly expressing TetX without shaking (F), with 10s shaking (G), with 50s shaking (H). Light grey is 0 µg/mL tetracycline, dark grey is 0.3125 µg/mL tetracycline, black is 0.625 µg/mL tetracycline. The red line represents a statistically significant biphasic curve as judged by a Bonferroni multiple comparisons test (treated with 0.625 µg/mL tetracycline). Points represent mean values, and error bars represent standard error of the mean. Mean and SEM are calculated using six biological replicates.

At 0 s/min shaking and 0.625 or 0.3125 µg/mL tetracycline, the relative growth of cells with the TetX module decreased (**Fig. 3B** black, dark gray, **Supp. Fig. 6A** 2^nd^ & 3^rd^ columns, **Supp. Fig. 7A** mid & bottom rows). With 0 s/min shaking and 0 µg/mL tetracycline, relative growth did not significantly change as gel concentration increased (**Fig. 3B** light gray, **Supp. Fig. 6A** 1^st^ column, **Supp. Fig. 7A** top row). At 10 s/min shaking and 0.625 µg/mL tetracycline, cells with the TetX module exhibited a significant biphasic change in relative growth as gel concentration increased (**Fig. 3C** red, **Supp. Fig. 6B** 3^rd^ column, **Supp. Fig. 7B** bottom row). With 10 s/min shaking and 0 or 0.3125 µg/mL tetracycline, the change in relative growth was visually biphasic, but had no significant trend (**Fig. 3C**, dark and light grey, **Supp. Fig. 6B** 1^st^ & 2^nd^ columns**, Supp. Fig. 7B** top and mid row). These results suggest that periodic perturbation interacts with TetX expression to allow bacteria to enhance growth in a manner sensitive to hydrogel concentration.

To distinguish whether biphasic growth was a general response to the presence of shaking or a unique response to a specific frequency of shaking, we gathered additional data from cells shaken at 50 s/min. At 50 s/min shaking and 0.3125 µg/mL tetracycline, relative growth decreased as hydrogel concentration increased (**Fig. 3D**, dark grey, **Supp. Fig. 6C** 2^nd^ column, **Supp. Fig. 7C** mid row). At 50 s/min shaking and 0 or 0.625 µg/mL tetracycline, relative cell growth did not significantly change (**Fig. 3D** black & light grey, **Supp. Fig. 6C** 1^st^ & 3^rd^ columns, **Supp. Fig. 7C** top & bottom rows). These results suggest the biphasic response of TetX-expressing cells grown with 10 s/min shaking and 0.625 µg/mL tetracycline (**Fig. 3C** red) is based on the frequency of periodic perturbation rather than the presence of perturbation.

To test whether the frequency-specific biphasic bacterial growth curve was mediated by the expression dynamics of the constitutive TetX module, we built and tested an inducible TetX module under the same shaking conditions and hydrogel concentrations. Inducible TetX expression was achieved by transforming the constitutive TetX module into a cell line (DH5*α*pro) that regulated module expression via genomically integrated *tetR* (**Fig. 3E**, **Supp. Fig. 8**, Methods-Section B). At 0 s/min shaking, no tetracycline concentrations produced a significant change in relative growth as hydrogel concentration rose (**Fig. 3F**, **Supp. Fig. 6D**, **Supp. Fig. 9A**). At 10 s/min shaking and 0.625 µg/mL tetracycline, the inducible TetX module displayed a significant biphasic relationship between relative growth and hydrogel concentration (**Fig. 3G** red, **Supp. Fig. 6E** 3^rd^ column, **Supp. Fig. 9B** bottom row). This biphasic growth curve of the inducible module was shifted to the right of the constitutive module growth curve, suggesting that the activation kinetics of TetX production affect population-level integration of spatial and temporal information. At 10 s/min shaking and 0 or 0.3125 µg/mL tetracycline, bacteria inducibly expressing TetX experienced no significant change in relative growth as hydrogel concentration rose (**Fig. 3G** light and dark grey, **Supp. Fig. 9B** top & middle rows). At 50 s/min shaking and 0, 0.3125, or 0.625 µg/mL tetracycline, the inducible TetX module similarly displayed no significant change in relative growth as hydrogel concentration increased (**Fig. 3H**, **Supp. Fig. 6D**, **Supp Fig. 9C**). Taken together, the results from the constitutive and inducible TetX modules (**Fig. 3**, **Supp. Fig. 6**, **Supp. Fig. 9**) demonstrate that the genetic regulation of an antibiotic resistance gene tunes the hydrogel sensitivity of bacterial growth under tetracycline exposure.

### A stochastic resonance model can explain the emergent growth dynamics of bacteria

After observing counterintuitive, frequency-sensitive bacterial growth, we sought to build a model to explain this phenomenon. Since protein expression level is similar between different shaking frequencies (**Fig. 2C**), differences in mean expression level are unlikely to drive biphasic growth with respect to hydrogel concentration. Alternatively, frequency-specific, biphasic responses have been previously associated with stochastic resonance in biological systems (McDonnell and Abbott, 2009; Paulsson et al., 2000; Zeng et al., 2009). For stochastic resonance theory to apply, there must be a weak oscillatory force and a source of biochemical noise (**Fig. 4A**).

**Figure 4.**
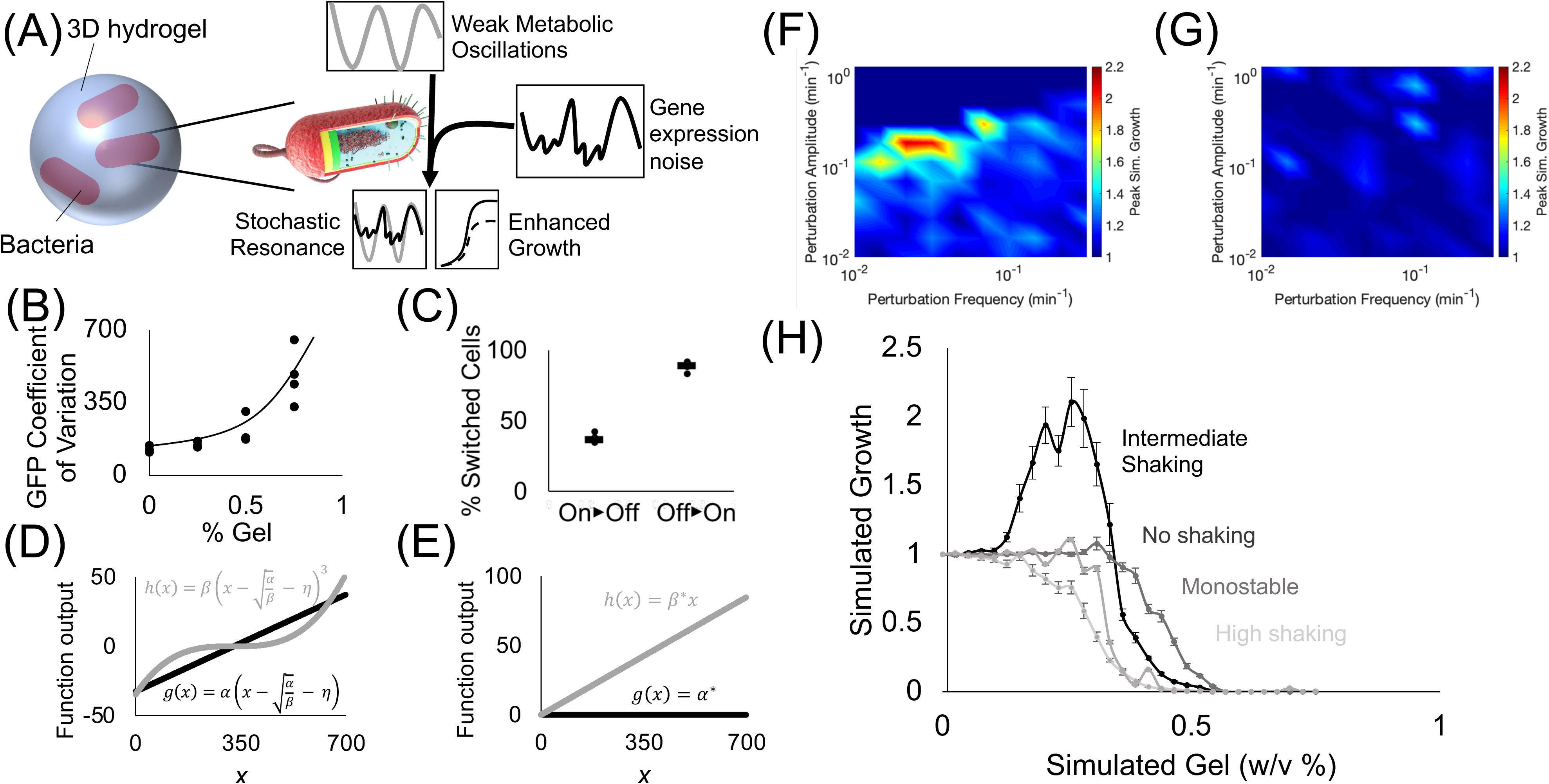
Simulated stochastic resonance generates biphasic, frequency-dependent bacterial growth. (A) Cartoon of the proposed model of bacterial stochastic resonance. At a specific frequency shaking, weak metabolic oscillations resonate with stochastic fluctuations in TetX production, boosting growth in tetracycline-treated bacteria. (B) Coefficient of variation of bacterial GFP expression in hydrogel. Graph shows that increased hydrogel concentration increases the variability of genetic module expression. Points represent individual replicates, line highlights trend. Four replicates per condition. Lines represent the median proportion of switched bacteria. (C) Quantification of ‘on’ and ‘off’ cellular populations that grew from cells sorted in the opposite state. Results suggest that gene expression can switch between ‘on’ and ‘off’ expression modes. Points represent individual replicates. Four replicates per condition. (D) Output of the positive (h(x), grey line) and negative function (g(x), black line) that represent a system with two stable fixed points. (E) Output of the positive (h(x), grey line) and negative function (g(x), black line) that represent a system with one stable fixed point. (F) Heat plot showing the simulated growth of bacteria at different frequencies and amplitudes of periodic perturbation. Color indicates the maximum growth of bacteria over a range of simulated hydrogel concentrations normalized by the growth of bacteria at the lowest gel concentration. Red areas show regions where stochastic resonance is causing biphasic changes in bacterial growth with respect to gel concentration. Surface plot values represent mean values. The mean is calculated using two hundred simulation replicates. (G) Heat plot showing the simulated growth of bacteria that cannot achieve bistable growth at different frequencies and amplitudes of periodic perturbation. Color indicates the maximum growth of bacteria over a range of simulated hydrogel concentrations normalized by the growth of bacteria at the lowest gel concentration. The results indicate that stochastic resonance does not occur under these conditions. Surface plot values represent mean values. The mean is calculated using two hundred simulation replicates. (H) Simulated bacterial growth in different hydrogel concentrations. Black line represents simulation with the amplitude, frequency, and bistable growth required for stochastic resonance. Black line shows growth under stochastic resonance conditions causes a biphasic relationship between the total growth in the system and the simulated gel concentration, similar to the 10 s/min shaking conditions in Fig. 3. Dark grey represents simulation with a periodic amplitude of zero, similar to the 0 s/min shaking conditions. Intermediate grey represents a simulation with a monostable system. Light grey represents a simulation with a reduction in the frequency of periodic perturbation. Results demonstrate that changing periodic forcing amplitude, frequency, or the underlying bistability of growth eliminates biphasic growth with respect to simulated gel concentration. Points represent mean values, and error bars represent standard error of the mean. Mean and SEM are calculated using two hundred simulation replicates.

The growth rate characterization experiments for Perceval show that there is an approximately 18% difference between the ATP/ADP ratio of cells with a significantly different growth rate (**Fig. 1B**). When cells are periodically perturbed, the maximum versus minimum A490/A405 measurements show the peak versus the trough of ATP/ADP oscillations are approximately 6% different (**Fig. 1C-D**). These results show that that shaking can exert an oscillatory force on bacterial metabolism that is not necessarily strong enough to be associated with a significant change in growth rate. This, in turn, suggests that periodic shaking exerts a weak oscillatory influence on growth.

Hydrogels have been hypothesized to generate extrinsic noise in encapsulated bacteria. Because real-time fluctuation of protein levels in single planktonic cells under shaking conditions cannot be measured, we used flow cytometry to infer the effect of hydrogel encapsulation on gene expression noise. Bacteria encapsulated in hydrogel experience an increase in the coefficient of variation of fluorescent reporter expression (**Fig. 4B**) caused by the splitting of the population between ‘on’ and ‘off’ expression modes (**Fig. 2B**). We tested the ability of cells grown in hydrogel to switch between ‘on’ and ‘off’ states by sorting the two populations, growing them independently, then quantifying changes in the proportion of ‘on’ vs. ‘off’ cells (Methods-Section H). A mixture of ‘on’ and ‘off’ states were observed in cultures grown from cells started from exclusively one state. These results suggest that cells dynamically switch between ‘on’ and ‘off’ states. These results build off prior work showing that synthetic materials can increase bacterial heterogeneity (Alvarez et al., 2009; Pabst et al., 2016; Wessel et al., 2013), demonstrating that cells grown in hydrogel exhibit dynamic fluctuations of gene expression (**Fig. 4C**). Based on these experiments, we assert that hydrogel causes dynamic fluctuations in TetX expression, imparting noise on the growth rate of encapsulated cells.

After establishing the biological criteria for stochastic resonance in synthetically encapsulated bacteria, we modified a classical stochastic differential equation (Eq. 1) (Gammaitoni et al., 1998) to describe the effect of stochastic resonance on bacterial growth (Methods-Section I).

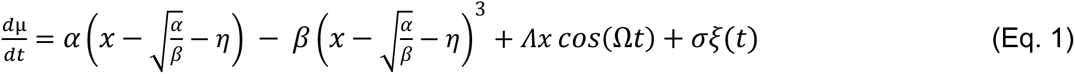

Where *µ* is the growth rate, *x* is the concentration of cells normalized by the maximum concentration of cells, *t* is time, *α* and *β* represent the growth of bacteria expressing the TetX module under tetracycline treatment (min^-1^), *η* represents the basal bacterial concentration normalized by the maximum concentration of cells, *Δ* is the amplitude of a weak periodic effect on growth (min^-1^), *Ω* is the period of a weak periodic effect on growth (min^-1^), and *σ* is the noise magnitude (min^-1^).

The bacterial growth term represented by *α*, *β*, and *η* was chosen to represent bistable growth caused by the feedback between importation, degradation, and translation-inhibition activity of antibiotics as described by Deris et. al. (Deris et al., 2013). To verify the bistability of the equation, we plotted the output of the positive and negative function terms to identify the stable and unstable fixed points (**Fig 4D**). We also implemented a simpler equation (Eq. 2) to represent non-bistable (i.e. monostable) growth of bacteria in the absence of tetracycline.

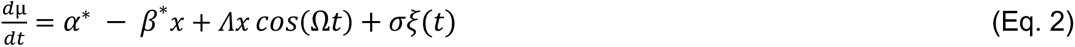

Where *µ* is the growth rate, *x* is the concentration of cells normalized by the maximum concentration of cells, *t* is time, *α*^∗^ and *β*^∗^ represent the growth of bacteria in the absence of tetracycline (min^-1^), *Δ* is the amplitude of a weak periodic effect on growth (min^-1^), *Ω* is the period of a weak periodic effect on growth (min^-1^), and *σ* is the noise magnitude (min^-1^). A parameter set for Eq. 2 was empirically derived to quantitatively match the integral of the growth rates from Eq. 1. The stability diagram of Eq. 2 was plotted under the chosen parameters, demonstrating that it is not bistable (**Fig. 4E**).

Next, Eq. 1 and 2 were used to simulate bacterial growth with weak periodic perturbation of growth rates. To capture the effect of hydrogel encapsulation on bacterial growth rate, noise was applied to the system such that it increased the heterogeneity of growth rates while also decreasing the overall growth rate. Simulations showed that at a narrow range of amplitude and frequency, the relative bacterial growth rate over a range of simulated gel concentrations was enhanced (**Fig. 4F**). Simulated bacterial growth without bistability did not experience frequency and noise specific enhancements (**Fig. 4G**).

The simulated growth of bacteria experiencing frequency-dependent enhancement was plotted against the range of simulated gel concentrations (**Fig. 4H** black line). Under conditions where relative growth enhancement was reported, simulated growth was found to be biphasic as simulated hydrogel concentrations increased (**Fig. 4H** black line). Removing the weak periodic force (**Fig. 4H** dark grey), changing the underlying bistability (**Fig. 4H** mid grey), or altering the periodic forcing frequency (**Fig. 4H** light grey) eliminated the biphasic growth with respect to simulated hydrogel concentration. These simulations demonstrate that stochastic resonance is sufficient to explain the biphasic, frequency-dependent bacterial growth curves that we experimentally observed in tetracycline-resistant bacteria.

## Discussion

Encapsulation of designer bacteria into hydrogels has birthed a new class of smart devices that can internally coordinate decision-making. These engineered living materials require novel strategies to maintain the potency of their biological components under restrictive metabolic conditions. To address this need, the effects of internal genetic machinery and external kinetic perturbation were systematically investigated in bacteria encapsulated in a three-dimensional hydrogel. An antibiotic resistance module was shown to optimize bacterial growth under restrictive encapsulation conditions based on the frequency of a rapid periodic disturbance. A simple biophysical model (Eq. 1) of stochastic resonance was proposed to explain this information processing phenomenon that appears to coordinate bacterial growth in three-dimensional hydrogels.

The model developed in this study provides a framework to maximize the efficiency of encapsulated bacterial growth by coordinating the timing of intra- and extra- cellular fluctuations. This model could be used as a guide for bacteria-seeded materials with gene expression noise tuned to resonate with a natural periodic process like peristalsis. Conversely, this model could also be used to determine which physical properties of an encapsulating substrate would be optimal to boost pre-existing cellular processes.

This study invites several lines of future inquiry. New strains and encapsulated substrates may be tested to create more systematic guidelines for the application of stochastic resonance to engineered living materials. To explain why constitutive TetX-producing cells have a distinct optimal hydrogel concentration from inducible TetX-producing cells, additional genetic circuits could be used to explore which regulatory features mediate the integration of spatial and temporal information. Given the frequency sensitivity of stochastic resonance, full confirmation of the model proposed in this study will require real-time tracking and measurement of single-cell gene expression in a semisolid environment. Further high-level evidence for stochastic resonance could also be collected by identifying a periodic shaking frequency that shifts the biphasic growth curve rather than eliminating it.

## Methods

### Contact for reagent and resource sharing

Further information and requests for resources and reagents should be directed to and will be fulfilled by the Lead Contact, Cheemeng Tan.

#### A. Media preparation and handling

All media in this study was prepared from granulated Miller’s Luria Broth from Research Products International. Seaplaque low-melting-temperature agarose (VWR) was incorporated into the media before autoclaving. Media was autoclaved for 30 minutes at 121°C. Sterilization and pressure release took one hour, after which media was cooled on the bench for 10 minutes before being placed in a 37°C incubator and being shaken at 200 rpm. During the course of experiments where cells were grown with low-melting-temperature-agarose, media was placed on a hotplate kept at 40°C while it was in use, then returned to the shaker when it was not in use. Media was made fresh the day before every experiment.

Consistent sterilization time and continuous heating of media containing low-melting temperature agarose were found to be critical for the reproducibility of all results. Concentrations of low-melting-temperature agarose above .75% increased the variability of the individual replicates such that reproducible results were not possible and were thus excluded from this study.

#### B. Plasmid construction and expression control

All plasmids used in this study were generated by Gibson cloning. The coding sequence for TetX was ordered from Genscript. The TetX sequence, the GFP sequence, all promoters, ribosome binding sites, and degradation tags were amplified by PCR with Q5 polymerase (NEB), gel extracted (Qiagen), and quantified using a Nanodrop (Thermo Fisher). For the constitutive and inducible circuits, TetX and GFP sequences were assembled with P_Tet_, RBS, and ssra tags as necessary in the pZ backbone that had been digested with *Eco*RI and *Bam*HI with buffer 3.1 (all NEB) at 37°C in a static incubator overnight. For the inducible circuits, expression was controlled by a gnomically integrated copy of TetR in the line DH5*α*pro. DNA fragments were assembled via the Gibson Assembly Master Mix (NEB) for 1 hour at 50°C following the DNA molar ratios suggested by the company’s protocol. Successful transformations were assayed by digestion with *Nco*I, *Bam*HI, and buffer 3.1 and confirmed by Sanger sequencing. The plasmid for the luminescence tests was pCS-P_EsaR_-P_lux_, a gift from Cynthia Collins (Addgene plasmid #47655). *E. coli* DH5α/pRsetB-his7-Perceval was a gift from Gary Yellen’s lab (Addgene plasmid #20336). Plasmid pRsetB-his7-Perceval was isolated and transferred into *E. coli* DH5α/pSC-Ptet-T7 which expresses T7 polymerase constitutively.

#### C. Perceval characterization of the effect of periodic shaking on cellular metabolism

A single colony of *E. coli* DH5α/pSC-P_tet_-T7/pRsetB-his7-Perceval was picked and inoculated into LB with 50 μg/mL kanamycin and 100μg/mL carbenicillin overnight at 37°C at 120° tilt from horizontal with 200 rpm shaking.

For quantification of ATP/ADP in M9 media, the overnight culture was subcultured into M9 with 0.05%, 0.1%, 0.2% and 0.4% glucose with the corresponding antibiotic with an initial OD_600 nm_ of 0.05. Next, 200 µL cell culture was put inside 96 well black plate with a clear flat bottom (Corning, ref #3631) combined with a clear lid. ATP/ADP ratios were measured by taking three rounds of reading at excitation 495 nm/emission 530 nm, and 405 nm/emission 530 nm by plate reader in a 2 mins cycle with 10 s orbital shaking (amplitude 3). T-tests were performed in Microsoft Excel using two tails and unequal variance.

For quantification of ATP/ADP in LB medium, the overnight culture was subcultured into fresh LB at 37°C at 120° tilt from horizontal with 200 rpm shaking with initial OD_600 nm_=0.05, A490 and A405 values of *E. coli* DH5α/pSC-P_tet_-T7/pRsetB-his7-Perceval in LB medium with glucose were recorded using plate reader. ATP/ADP ratios were measured by taking three rounds of reading at excitation 495 nm/emission 530 nm, and 405 nm/emission 530 nm by plate reader in a 2 mins cycle with 10 s orbital shaking (amplitude 3). Subsequent readings were analyzed by Fourier transform in MATLAB using the fft function. Significance between differences in fluctuation amplitude was measured by a two-tailed, unequal variance t-test in Microsoft Excel.

#### D. Luminescent characterization of the effect of periodic shaking on cellular metabolism

All luminescence results were generated using a plate reader (Tecan M1000pro). Cells were grown overnight at 37°C at 120° tilt from horizontal with 200 rpm shaking in LB with antibiotics from a single colony, then diluted to a constant OD600 value of 0.5. All cell lines were incubated on ice for 30 minutes. Cell cultures standardized in this way were further diluted 1:100 into LB media with the appropriate antibiotics and the appropriate concentration of low-melting-temperature agarose as indicated in the data. DH5α cells were grown in media without antibiotics, the luminescent cell line was grown with 30 µg/mL kanamycin. These cells were incubated for 1.5 hours at 37°C at 120° tilt from horizontal with 200 rpm shaking until they reached the beginning of exponential growth, at which point they were distributed into a 96-well plate, covered with a lid, and put into the plate reader at 37°C. Total well volume was 200 µL. The plate reader subjected the cells to 10 seconds of orbital shaking (amplitude 3), iterated through three rounds of reading luminescence then OD600, then restarted the cycle. Each cycle took two minutes. Readings were taken for at least six hours. Comparisons between shaking and non-shaking conditions were made by picking times points where the OD600 readings of the shaking and non-shaking cultures were equal to 0.25.

#### E. Flow cytometry for cells expressing GFP

DH5α cells containing the pZ(P_Const_-GFP) construct were grown overnight at 37°C at 120° tilt from horizontal with 200 rpm shaking in LB with 30 µg/mL kanamycin from a single colony, then diluted to a constant OD600 value of 0.5. Cells were incubated on ice for 30 minutes, then further diluted 1:100 into LB media with 30 µg/mL kanamycin and the appropriate concentration of low-melting-temperature agarose as indicated in the data. These cells were incubated for 1.5 hours at 37°C at 120° tilt from horizontal with 200 rpm shaking until they reached the beginning of exponential growth, at which point they were distributed into a 96-well plate, covered with a lid, and put into the plate reader at 37°C. Total well volume was 200 µL. OD600 and GFP fluorescent intensity of each well was monitored in the plate reader with measurements taken every 10 minutes, with 10 seconds of orbital shaking (amplitude 3) every minute. Cells were grown in the plate for 2 hours before being diluted 1:100 into PBS with .4% paraformaldehyde. Fixed cells were run on a Novocyte Flow Cytometer (ACEA Biosciences). At least nineteen thousand events were collected for each sample, and data from each sample was analyzed using FCS Express version 4.0 (De Novo Software, Los Angeles, CA).

#### F. Characterization of biochemical circuits and relative growth

All growth results were generated using a plate reader (Tecan M1000pro). Cells were grown overnight at 37°C at 120° tilt from horizontal with 200 rpm shaking in LB with antibiotics from a single colony, then diluted to a constant OD600 value of 0.5. All cell lines were incubated on ice for 30 minutes to ensure uniform growth between different cell lines. Cell cultures standardized in this way were further diluted 1:100 into LB media with the appropriate antibiotics and the appropriate concentration of low-melting-temperature agarose as indicated in the data. The non-resistant cell lines, DH5α and DH5αpro, were grown in media without antibiotics, both constitutive lines and the inducible lines were grown with 30 µg/mL kanamycin, and the oscillator TetX lines were grown with 30 µg/mL kanamycin and 50 µg/mL carbenicillin. These cells were incubated for 1.5 hours at 37°C at 120° tilt from horizontal with 200 rpm shaking until they reached the beginning of exponential growth, at which point they were distributed into a 96-well plate, covered with a lid, and put into the plate reader at 37°C. Total well volume was 200 µL. OD600 and GFP fluorescent intensity of each well was monitored in the plate reader with measurements taken every 10 minutes. Without shaking, the plate was left undisturbed between measurements. Under intermediate shaking conditions, the plate experienced 10 seconds of orbital shaking (amplitude 3) every minute. Under high shaking conditions, the plate experienced 50 seconds of orbital shaking (amplitude 3) every minute. Cells were allowed to grow for 2 hours before 50 µL of culture per well was transferred into a new 96-well plate with 150 µL of media already in the plate along with indicated concentrations of tetracycline. Unless otherwise noted, all results were taken from cells after 120 minutes of growth in the second plate. Unless otherwise noted, all growth results were replicated 6 times: three sets of duplicated samples were collected on distinct days. No blinding or randomization was performed. A sample size of six was selected according to standard synthetic biology practices of sample replication inside a plate reader for significance testing, but these tests have no established effect size.

#### G. Significance testing on relative growth curves

Since we are trying to determine the significance of differences between many points in a gene expression transfer function, we chose to use a multiple comparisons test instead of a normal t-test. For experimental data, each relative growth curve (plotted with respect to noise intensity) was subjected to a Bonferroni multiple comparisons test, which performs significance testing between each point within the transfer function, correcting for false positives by adjusting the *p*-value of the test by dividing it by the number of comparisons being made. With this technique, we established confidence intervals between the values in each relative growth curve using a base *p*-value of 0.05. If the curve experienced a significant increase in cell growth followed by a significant decrease, that curve was classified as biphasic.

#### H. Measurement of stochastic switching expression in noisy environments

In brief, we followed the distribution of cells that started unimodally “on” and “off”, then used flow cytometry to characterize the bimodal populations of cells that were present at the end of the experiment and counted the proportion of the final population that had switched states. This experiment was broken into two phases to account for the competitive advantage of “on” cells under tetracycline treatment. The first phase starts at a high density of constitutively active cells before measuring their fluorescent gene expression while growing in 0.5% low-melting-temperature agarose. The second phase starts with a low density of cells sorted to be genetically inactive before measuring their fluorescent gene expression while growing in 0.5% low-melting-temperature agarose.

To measure the switching of cells from an “on” state to an “off” state, DH5α cells containing pZ(P_Const_-TetX-GFP) were grown overnight at 37°C at 120° tilt from horizontal with 200 rpm shaking in LB with 30 µg/mL kanamycin from a single colony, then diluted to a constant OD600 value of 0.5. Cells were incubated on ice for 30 minutes, then further diluted 1:100 into LB media with 30 µg/mL kanamycin and 0.5% low-melting-temperature agarose as indicated in the data. Total well volume was 200 µL. These cells were incubated for 1.5 hours at 37°C at 120° tilt from horizontal with 200 rpm shaking until they reached the beginning of exponential growth, at which point they were distributed into a 96-well plate, covered with a lid, and put into the plate reader with 0.625 µg/mL tetracycline at 37°C. OD600 and GFP fluorescent intensity of each well was monitored in the plate reader with measurements taken every 10 minutes, with 10 seconds of orbital shaking (amplitude 3) every minute. Cells were grown in the plate for 1 hour before being diluted 1:100 into PBS. Some of the cells were analyzed using a Novocyte Flow Cytometer (ACEA Biosciences), and some were used for cell sorting for the second phase of the experiment.

To measure the switching of cells from an “off” state to an “on” state, cells were sorted in an Astrios Cell Sorter (Beckman Coulter), defining the “off” population with a combination of gates on FITC height and width, thresholding reads on side scatter. Once sorted, 2500 of the “off” cells were put in LB media with 30 µg/mL kanamycin, 0.625 µg/mL tetracycline, and 0.5% low-melting-temperature agarose for a total well volume of 200 µL. The plate was covered with a plastic lid and grown for ∼ 12 hours at 37°C. OD600 and GFP fluorescent intensity of each well was monitored in the plate reader with measurements taken every 10 minutes, with 10 seconds of orbital shaking (amplitude 3) every minute. Once the cells had reached the same cell density that they had achieved in the first hour of growth, pZ(P_Const_-TetX-GFP)-containing cells were diluted 1:100 in PBS and analyzed using a Novocyte Flow Cytometer (ACEA Biosciences).

Data from each sample was analyzed using FCS Express version 4.0 (De Novo Software, Los Angeles, CA). Results were gated by only including counts that had a linear correlation between area and height measurement on the FITC fluorescent channel. The “on” population and the “off” population were separately determined to separate the two modes seen in the data, and these boundaries were uniformly applied to each replicate. For the “off” to “on” measurement, between five thousand and eighteen thousand counts were collected after gating. For the “on” to “off” measurement, between seventeen thousand and thirty-seven thousand counts were collected after gating.

#### I. Simulation of stochastic resonance using a stochastic differential equation

Simulation was performed in MATLAB. Equations were entered using the structures represented in Eq.1 or Eq. 2. The stochastic differential equation was implemented using the Euler-Maruyama method. 1000 simulations were done for each condition and averaged together. Average cell growth of each curve with respect to increasing noise was normalized to average cell growth at the lowest noise intensity.

## Data and software availability

The datasets and simulations generated during and/or analyzed during the current study are available from the corresponding author upon request.

## Acknowledgements

The work is supported by Human Frontier Science Program RGY0080-2015 (C.T.) and National Science Foundation CHE1808237 (C.T.). Research reported in this publication was supported by the National Institutes of Health under award number S10OD018223.

## Author contributions

DL and CT designed the study and wrote the manuscript. TG assisted with the characterization of the constitutive TetX construct, performed the Perceval experiments and the qPCR experiments. YX assisted with the characterization the gel effect on cell growth and the assembly of constructs used in this study.

## Supplementary Figure Legends

**Supplementary Figure 1.**
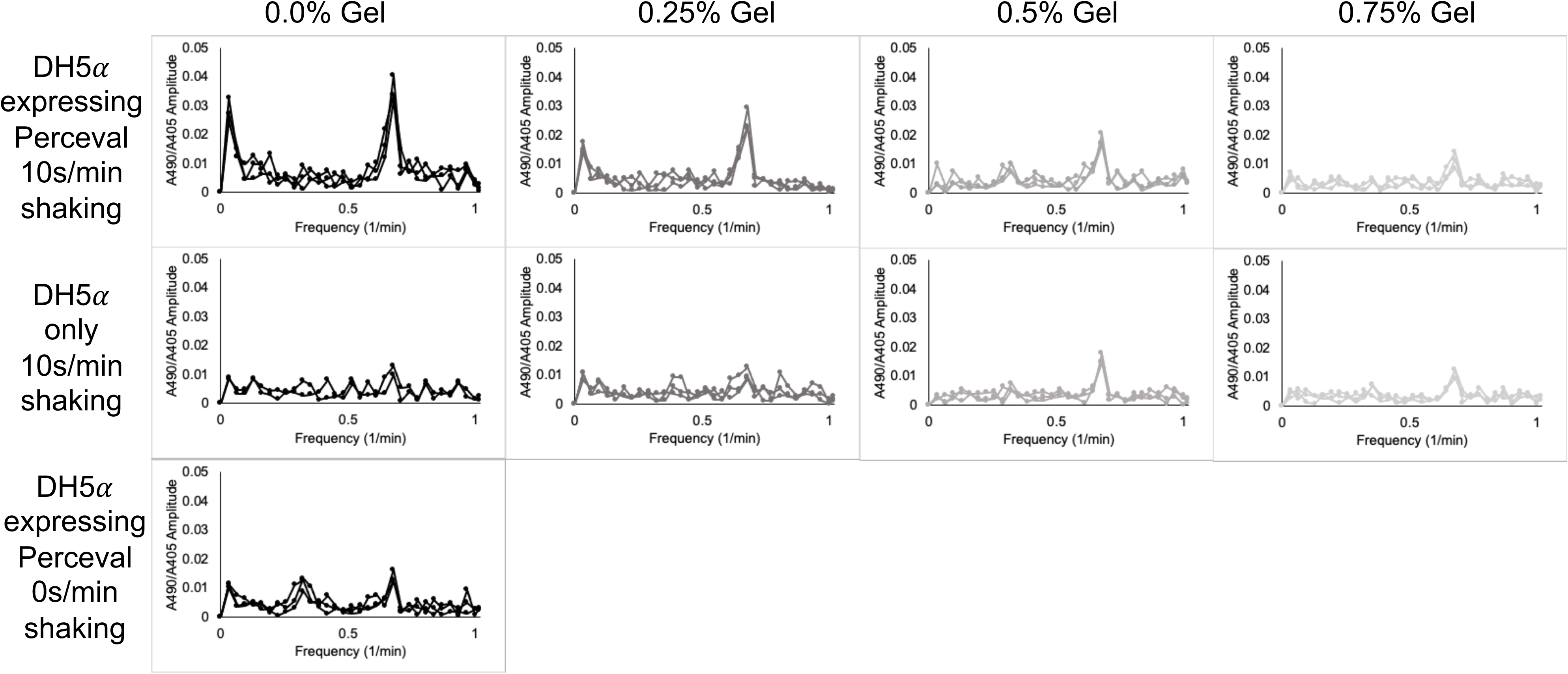
Frequency spectra of cells for A490/A405 ratio. Graphs representing Fourier transformations of the oscillating ratio of absorbance at 490nm and absorbance at 405nm. The ratio of A490/A405 has been shown to represent the ratio of ATP to ADP in cells expressing Perceval. These results show that the frequency spectra of cells with Perceval and cells without Perceval can be distinguished at 0.0% and 0.25% gel, but not 0.5% or 0.75% gel. Furthermore, the frequency spectra of cells expressing Perceval and shaken at 10s/min can be distinguished from cells expressing Perceval and shaken at 0s/min. All graphs show 3 biological replicates, except for DH5*α* cells in 0.0% gel with 10s/min shaking, which shows 2 biological replicates.

**Supplementary Figure 2.**
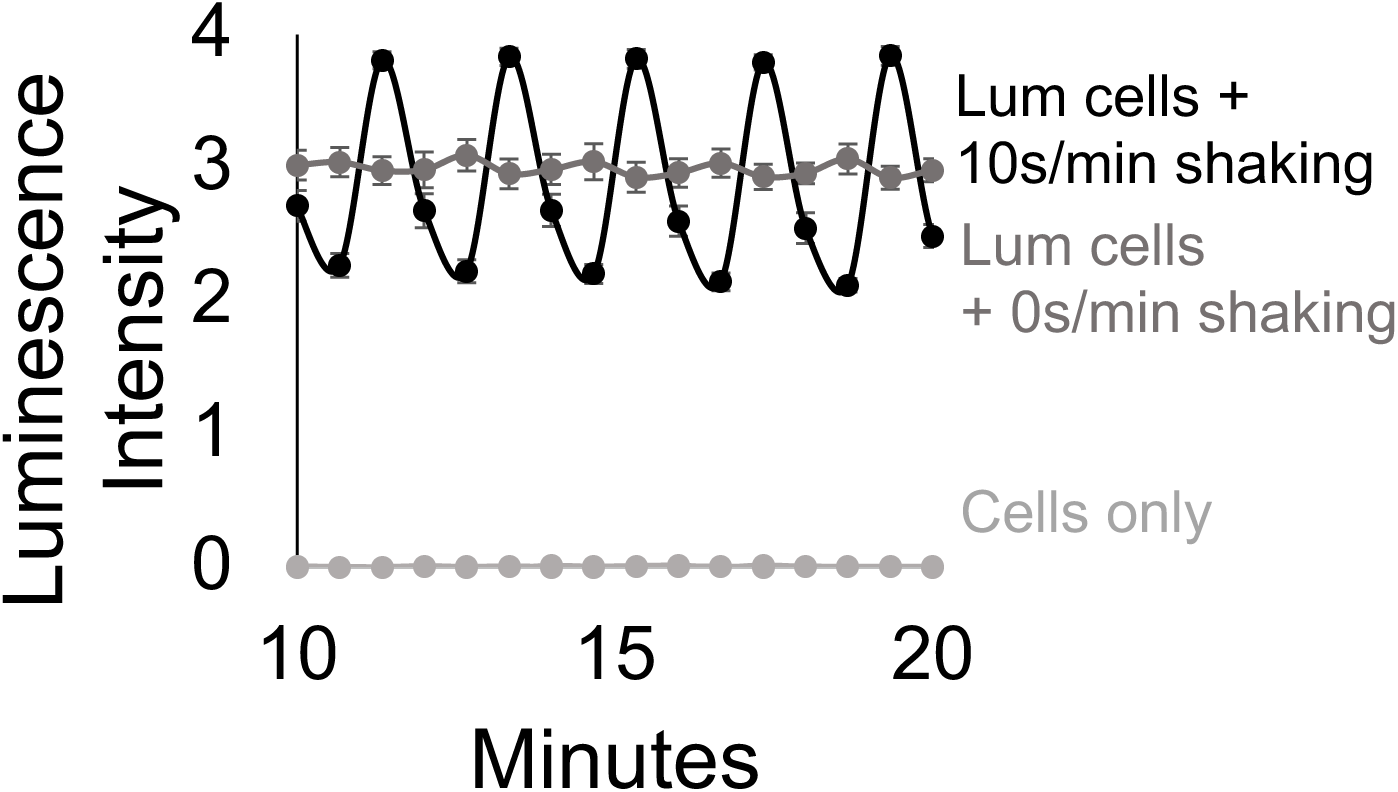
Bacteria expressing a luminescent reporter under periodic and heterogeneous conditions. Metabolic activity of cells in media with .5% low-melting-temperature agarose as measured by cells expressing luciferase, which uses an ATP- and oxygen-dependent reaction to catalyze the production of light. Periodic shaking induces an approximately two-fold periodic change in luminescence (Methods-section D). Along with fluctuations in Perceval A490/A405 levels, these results suggest that periodic shaking of cells imparts periodic perturbation of metabolism in those cells. Points represent mean values, and error bars represent standard error of the mean. Mean and SEM are calculated using six biological replicates.

**Supplementary Figure 3.**
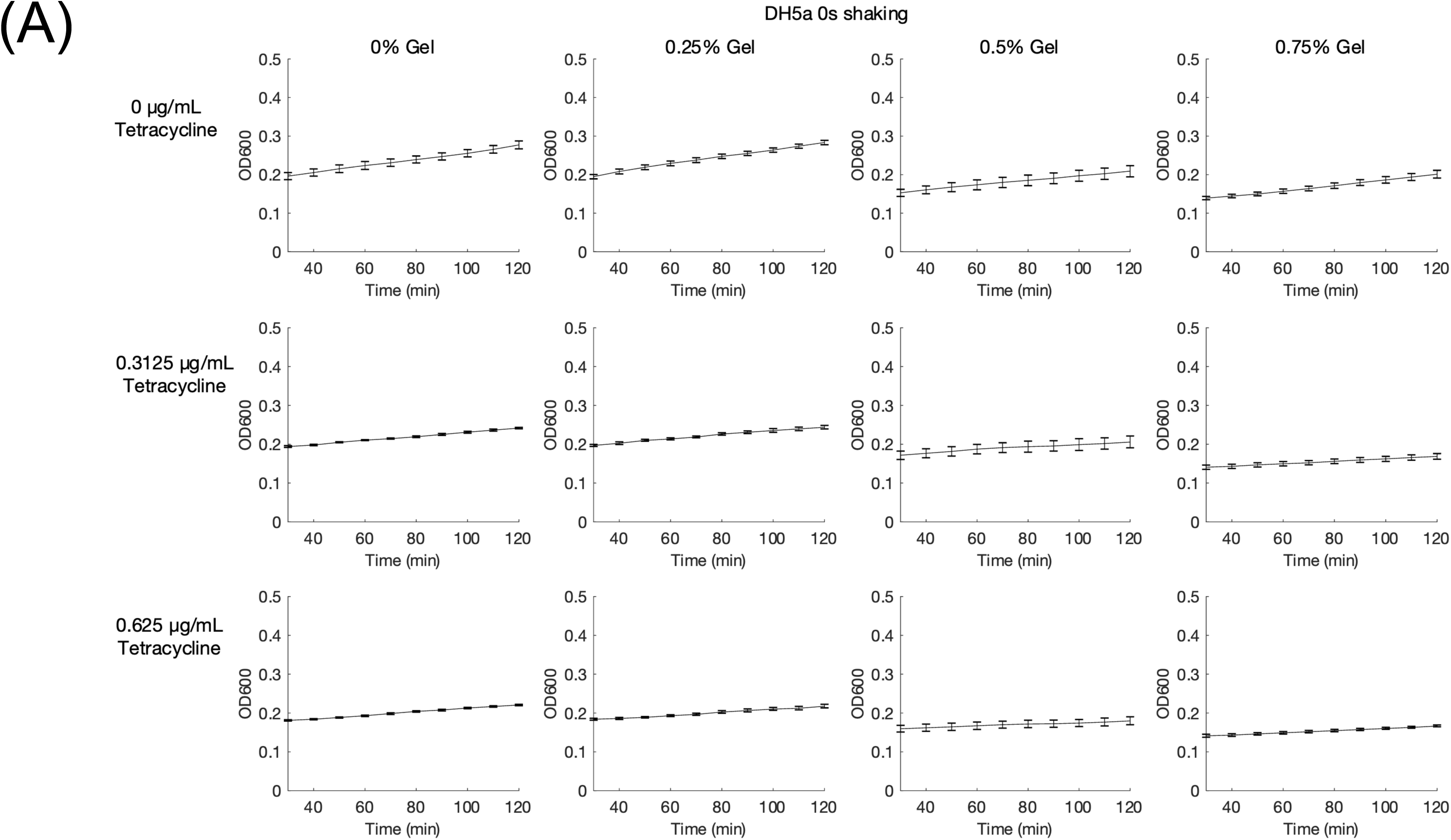
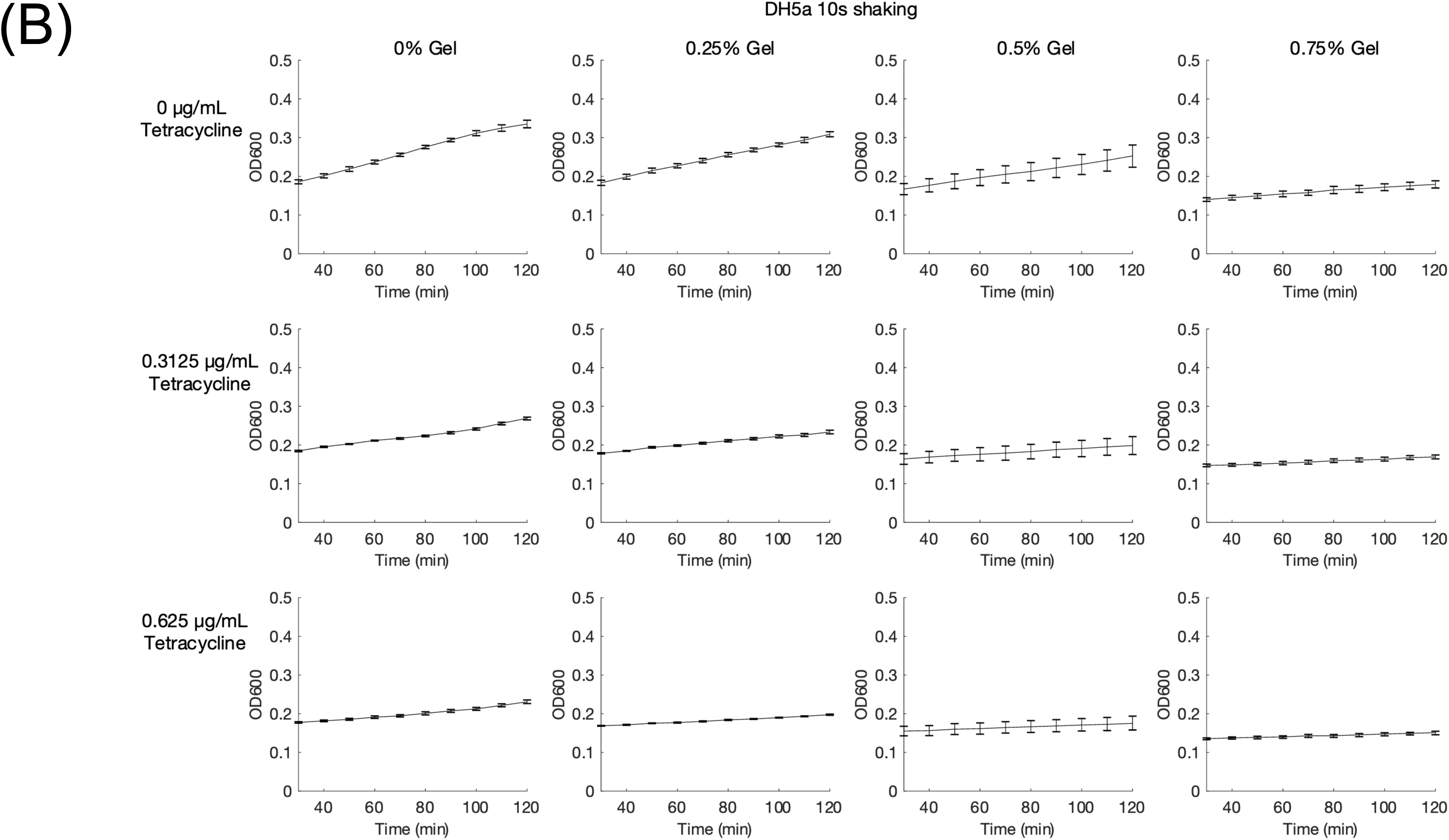
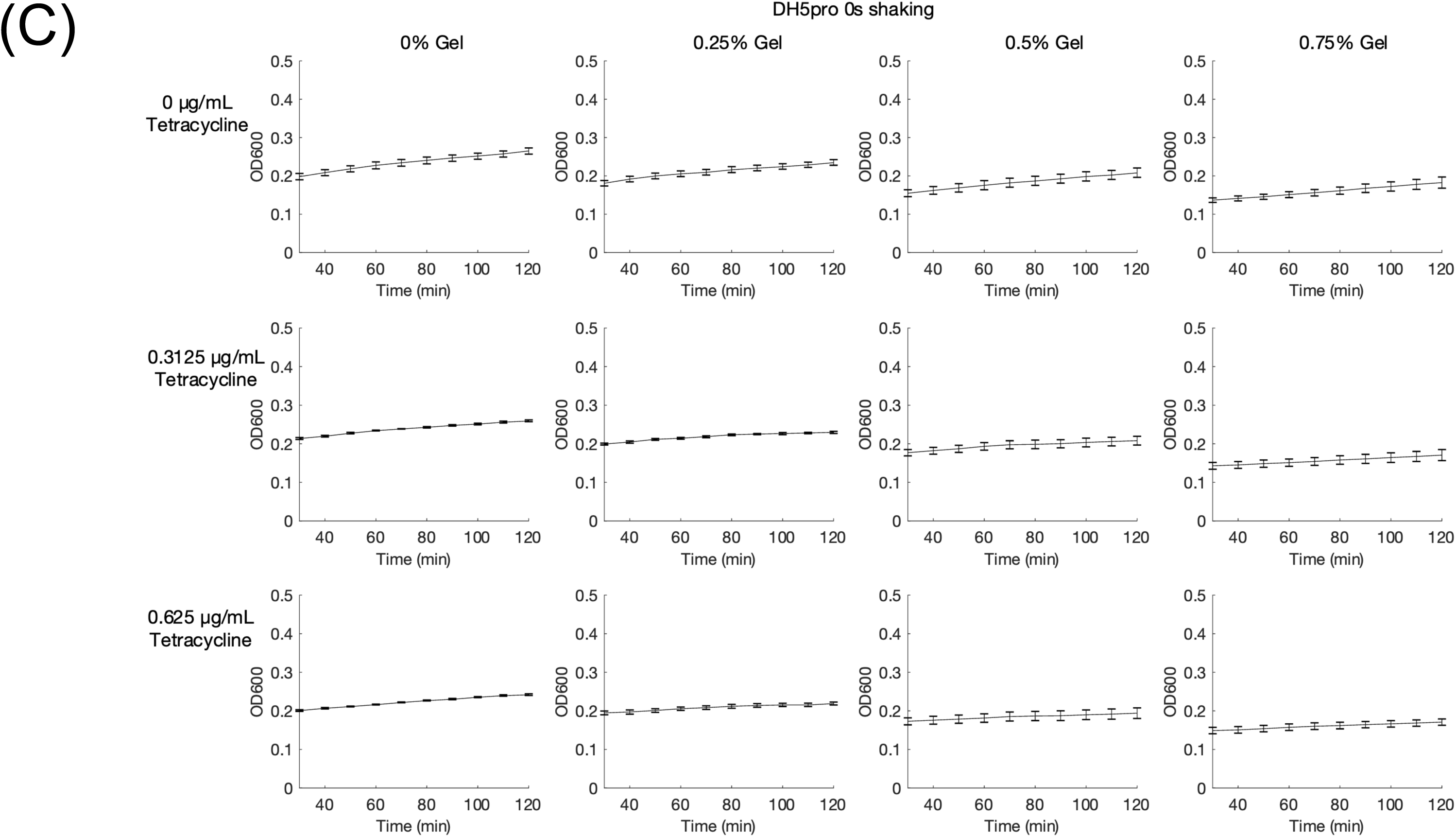
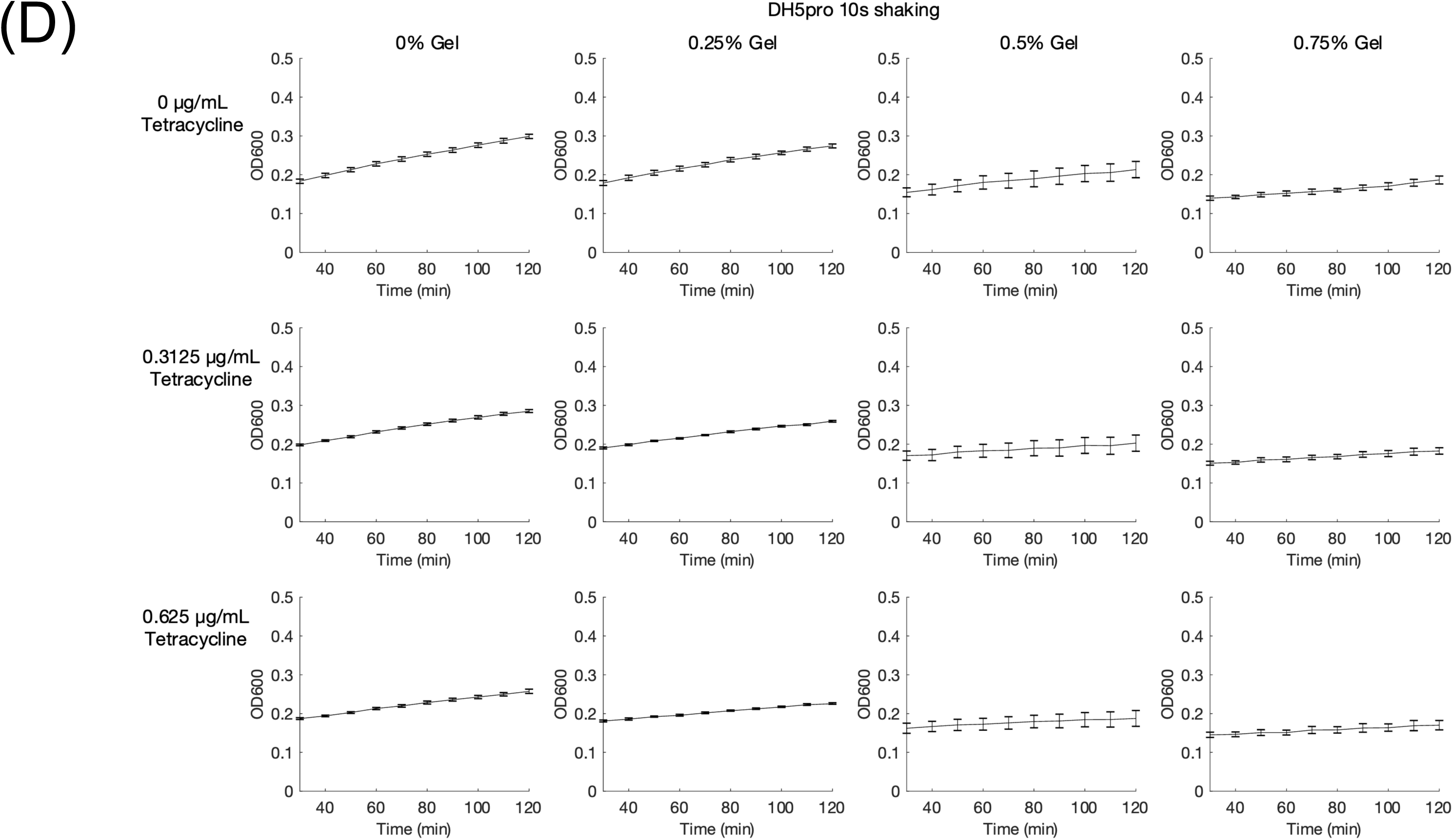
Changes in OD600 over time for DH5*α* and DH5*α*pro cells grown in hydrogel and tetracycline. (A) Changes in OD600 over time for DH5*α* grown without shaking. (B) Changes in OD600 over time for DH5*α* grown with 10 seconds shaking per minute. (C) Changes in OD600 over time for DH5*α*pro grown without shaking. (D) Changes in OD600 over time for DH5*α*pro grown with 10 seconds shaking per minute. (A-D) Cell grown in LB with 0, 0.25, 0.5, or 0.75% gel along with 0 µg/mL, 0.3125 µg/mL, or 0.625 µg/mL tetracycline. Points represent mean values, and error bars represent standard error of the mean. Mean and SEM are calculated using six biological replicates.

**Supplementary Figure 4.**
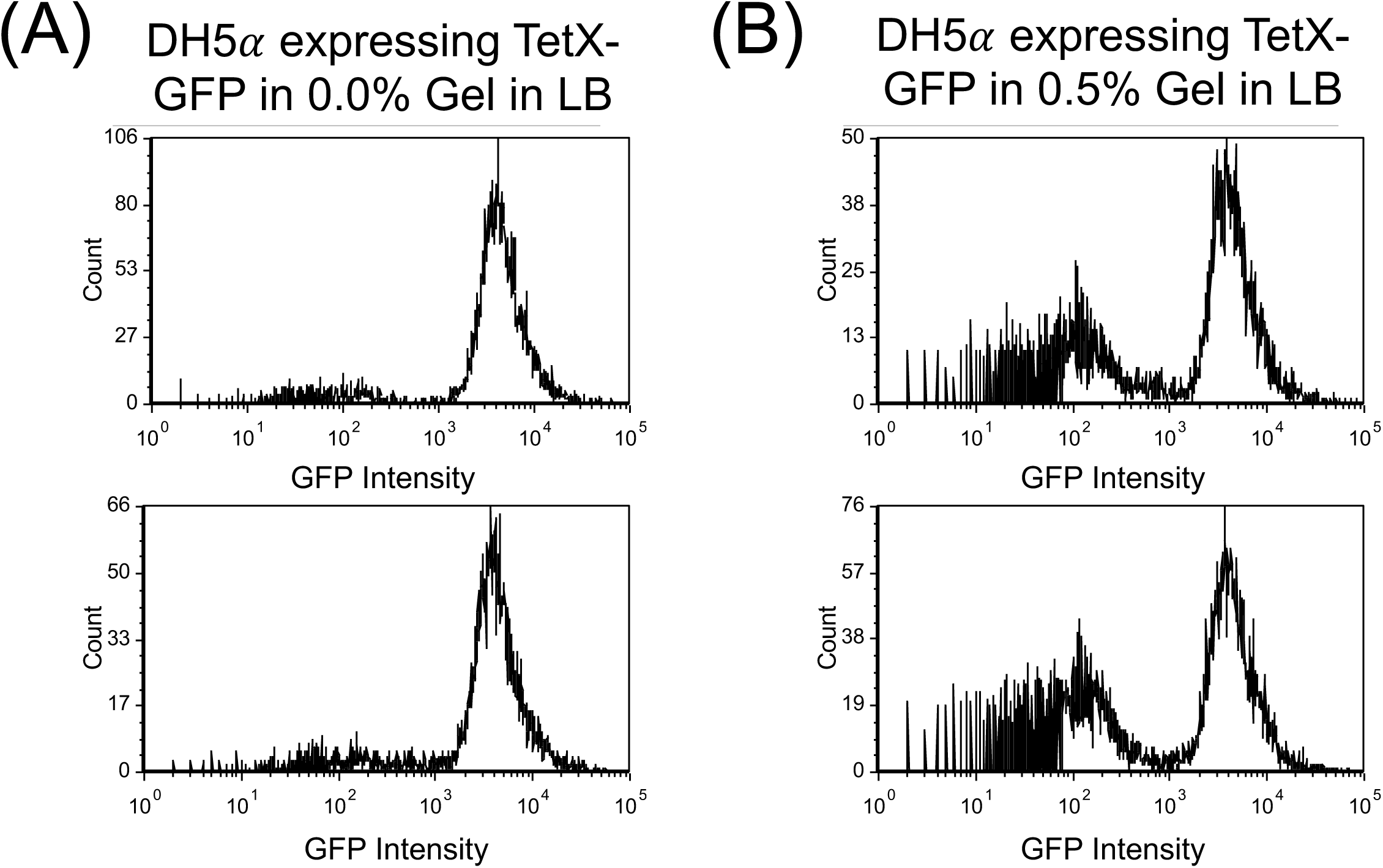
Hydrogel encapsulation causes heterogeneous protein expression. (A) Histograms of GFP intensity of DH5*α* cells constitutively expressing TetX-GFP in LB with 0% gel. All biological replicates are shown. (B) Histograms of GFP intensity of DH5*α* cells constitutively expressing TetX-GFP in LB with 0.5% gel. All biological replicates are shown. (A&B) Histograms represent 5000-8000 events after filtering.

**Supplementary Figure 5:**
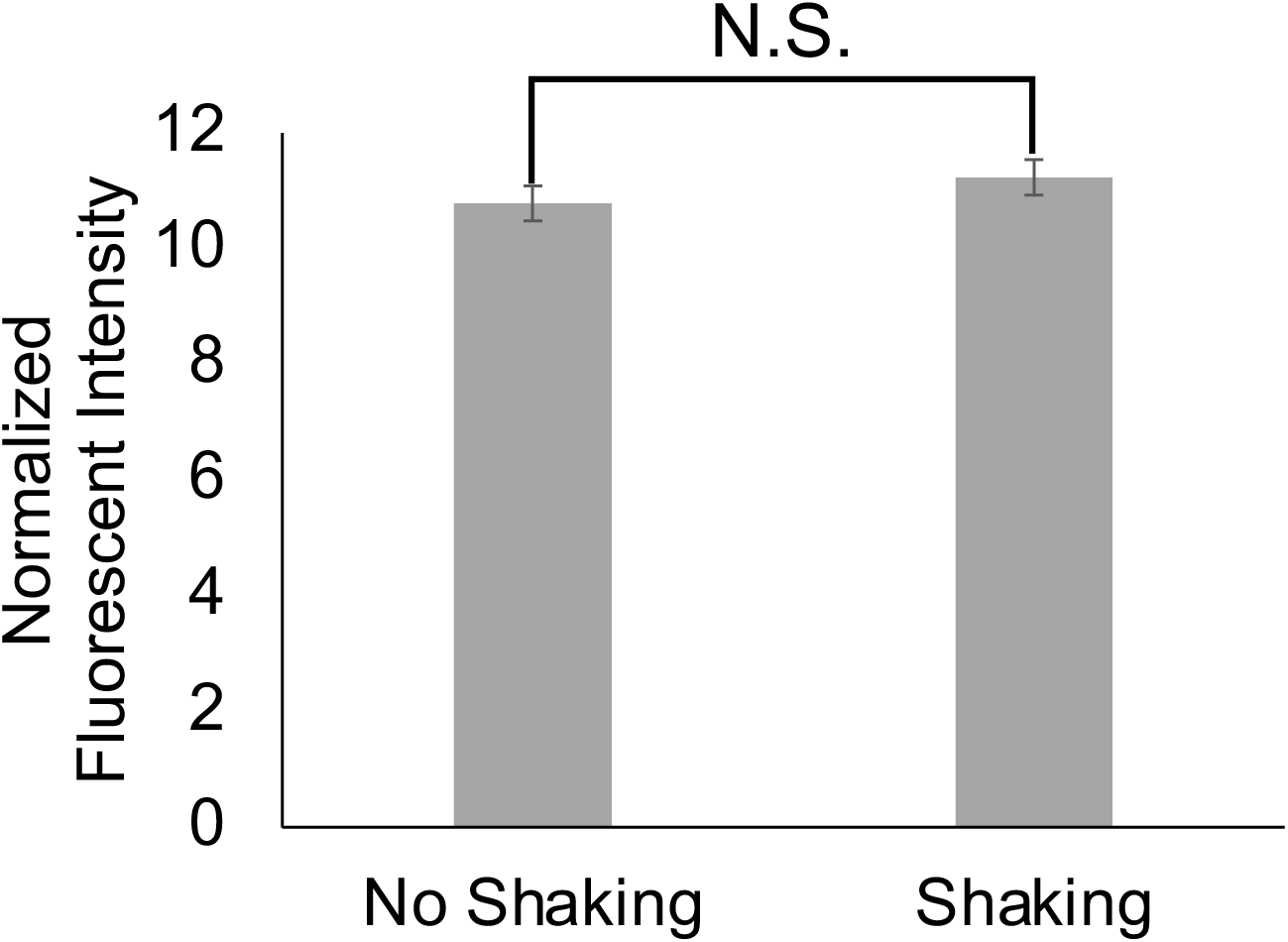
Periodic perturbation does not increase gene module expression. Fluorescent intensity of TetX-GFP normalized by OD600 of cells after 1 hour of growth. Two-tailed, equal variance t-test returns *p*-value = 0.345 N.S. stands for not significant. Bar height represents mean values, and error bars represent standard error of the mean. Mean and SEM are calculated using eight biological replicates.

**Supplementary Figure 6.**
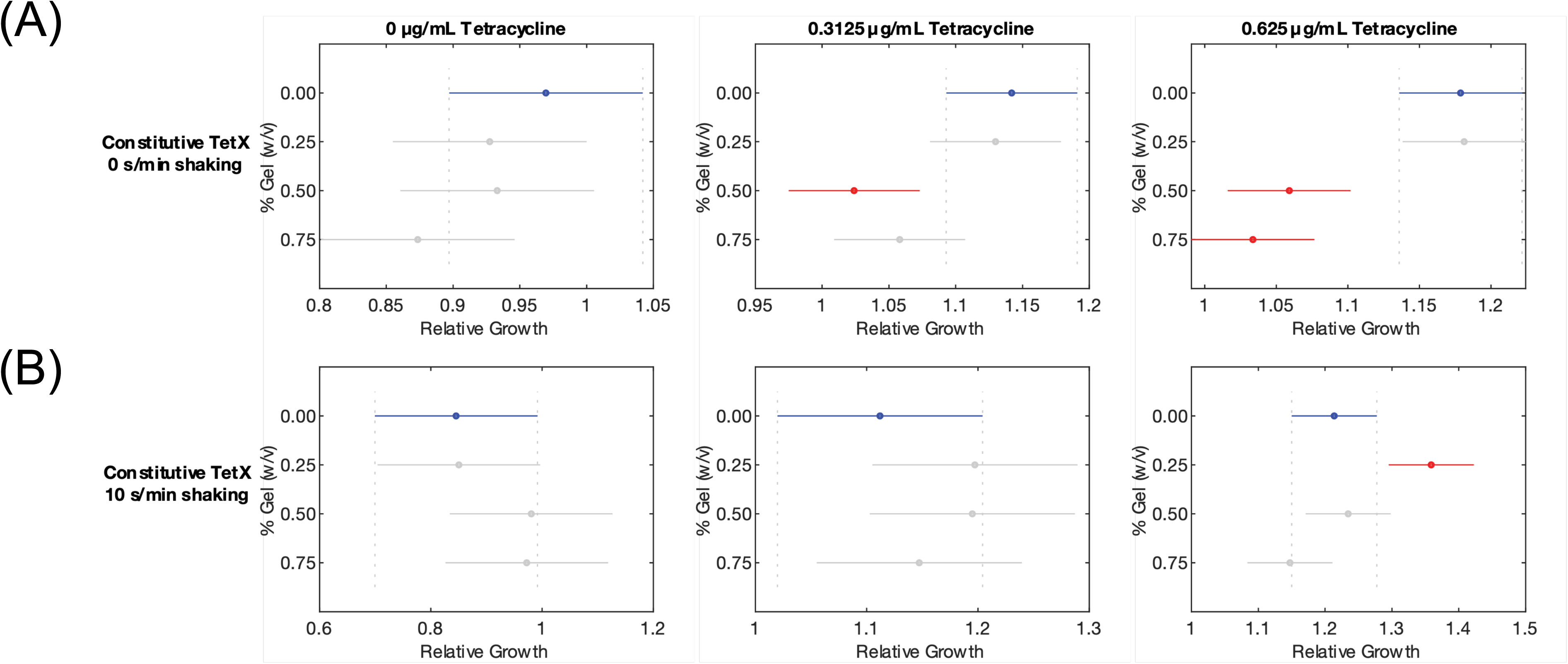
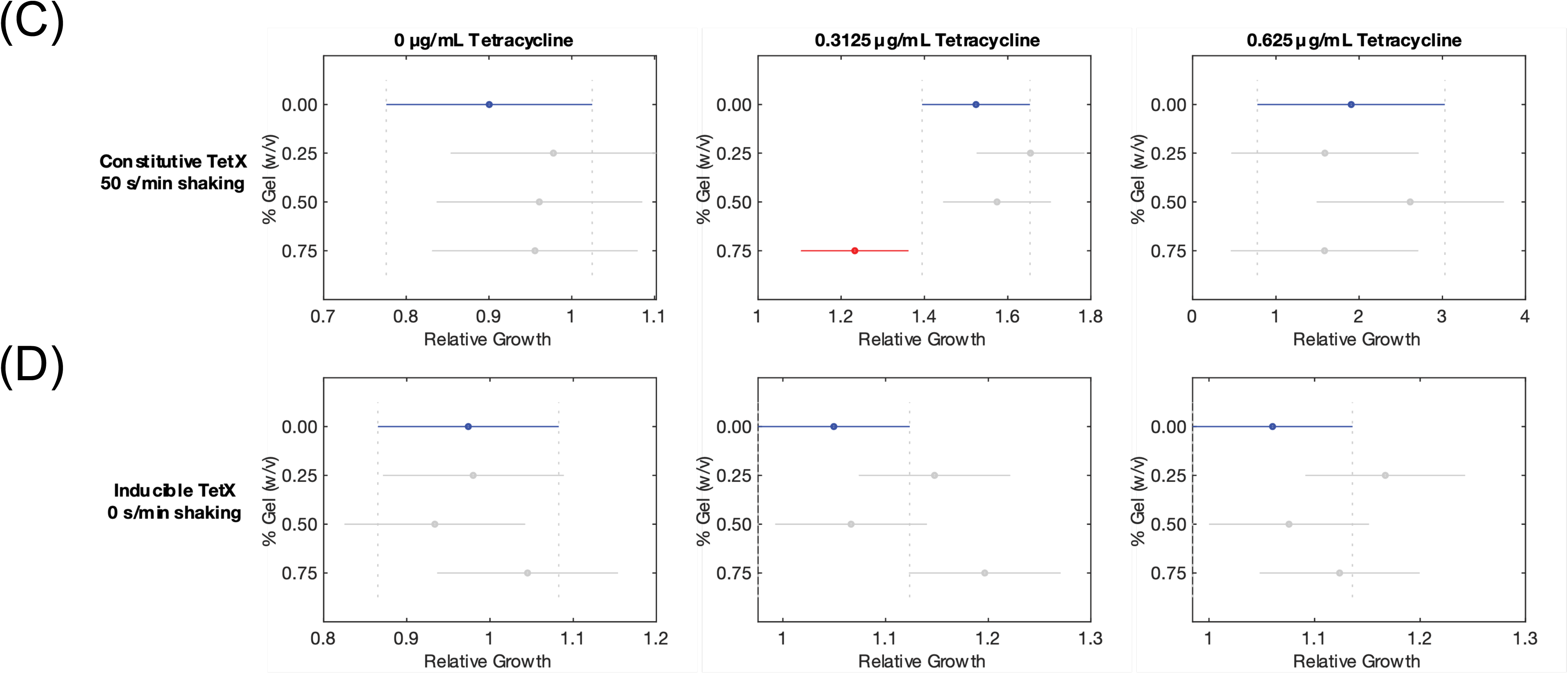
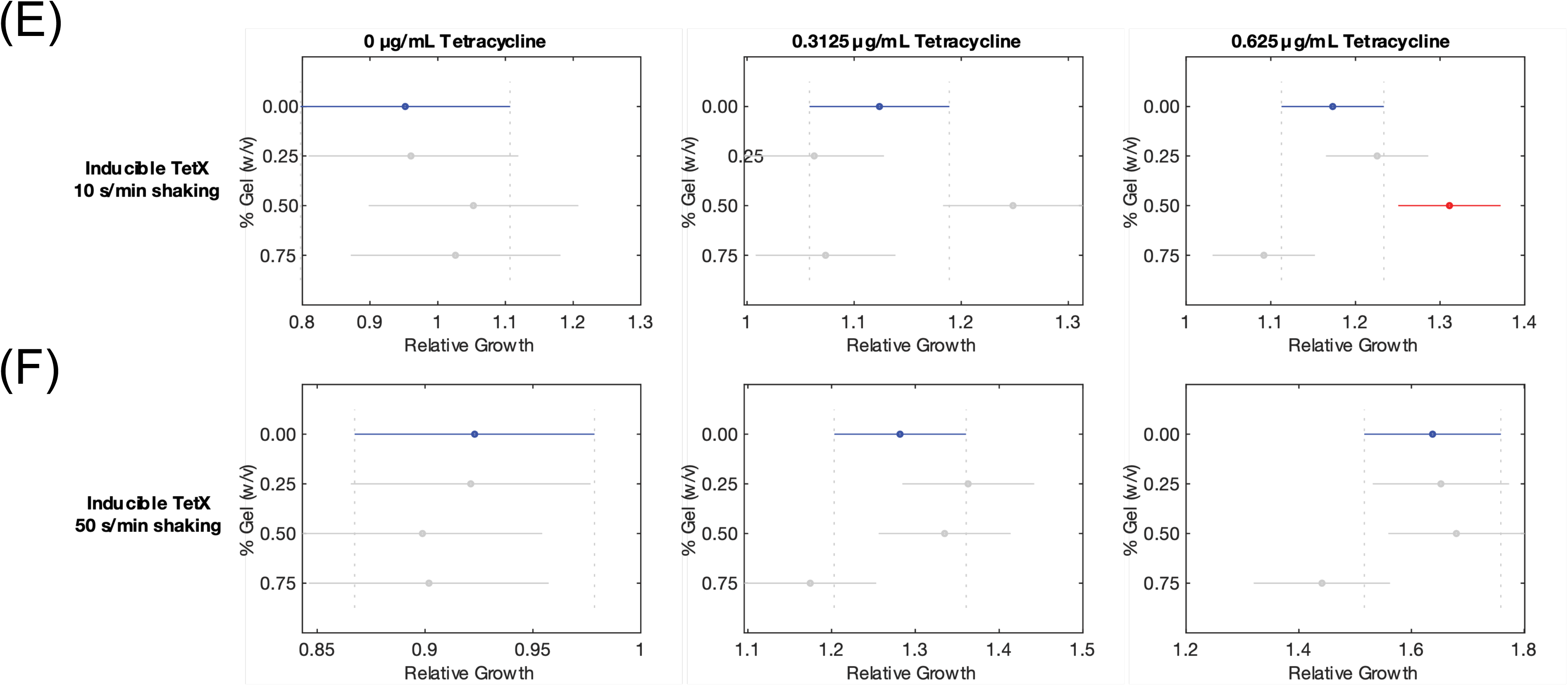
Significance testing of relative growth curves. (A) Confidence intervals of relative growth for bacteria constitutively expressing stable TetX grown with 0 s/min shaking (left panel) 0 µg/mL tetracycline (middle panel) 0.3125 µg/mL tetracycline (right panel) 0.625 µg/mL tetracycline (B) Confidence intervals of relative growth for bacteria constitutively expressing stable TetX grown with 10 s/min shaking (left panel) 0 µg/mL tetracycline (middle panel) 0.3125 µg/mL tetracycline (right panel) 0.625 µg/mL tetracycline (C) Confidence intervals of relative growth for bacteria constitutively expressing stable TetX grown with 50 s/min shaking (left panel) 0 µg/mL tetracycline (middle panel) 0.3125 µg/mL tetracycline (right panel) 0.625 µg/mL tetracycline (D) Confidence intervals of relative growth for bacteria inducibly expressing stable TetX grown with 0 s/min shaking (left panel) 0 µg/mL tetracycline (middle panel) 0.3125 µg/mL tetracycline (right panel) 0.625 µg/mL tetracycline (E) Confidence intervals of relative growth for bacteria inducibly expressing stable TetX grown with 10 s/min shaking (left panel) 0 µg/mL tetracycline (middle panel) 0.3125 µg/mL tetracycline (right panel) 0.625 µg/mL tetracycline (F) Confidence intervals of relative growth for bacteria inducibly expressing stable TetX grown with 50 s/min shaking (left panel) 0 µg/mL tetracycline (middle panel) 0.3125 µg/mL tetracycline (right panel) 0.625 µg/mL tetracycline (A-F) Significance intervals calculated by Bonferroni multiple comparisons test within the growth curve (Methods-Section G).

**Supplementary Figure 7.**
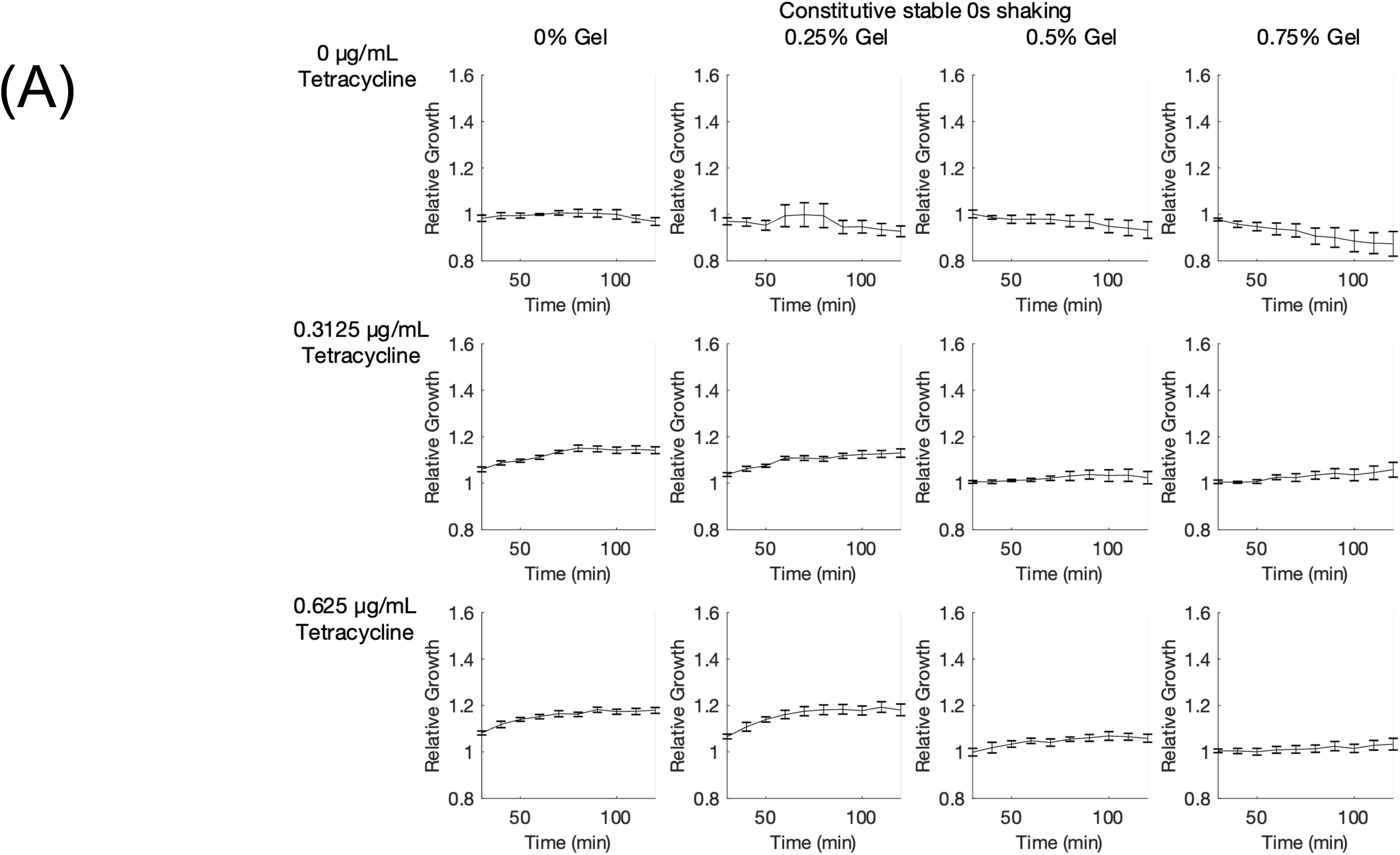
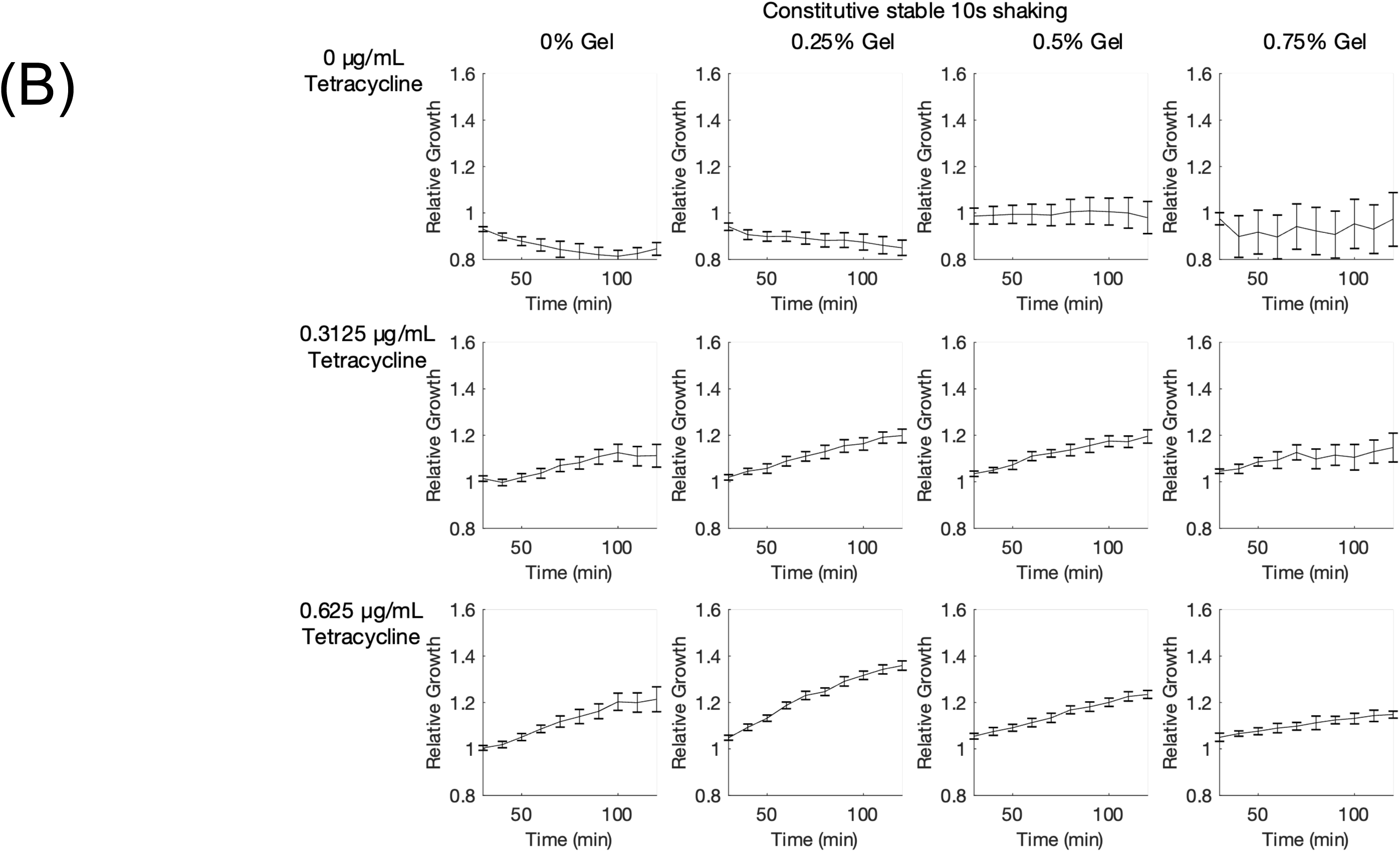
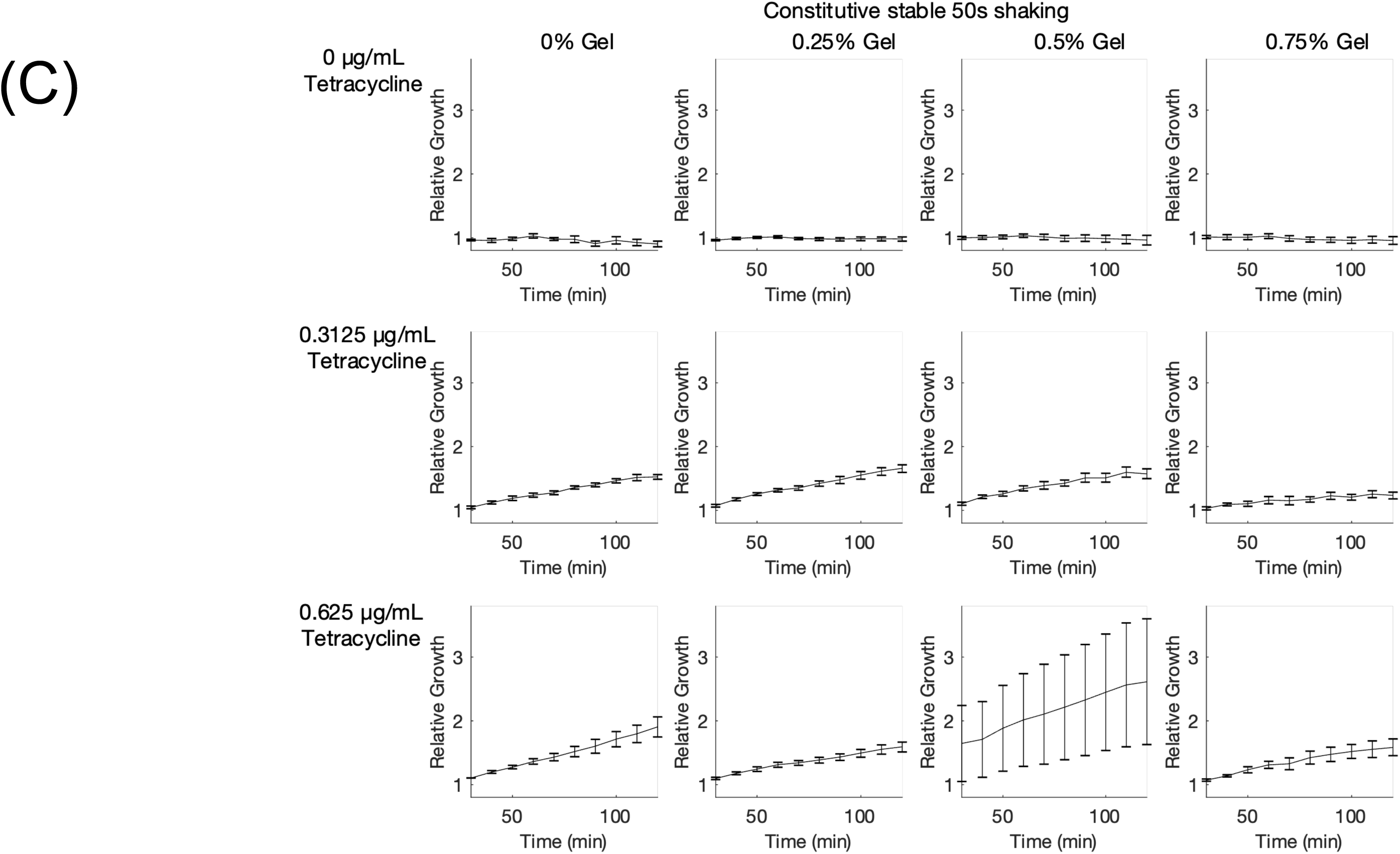
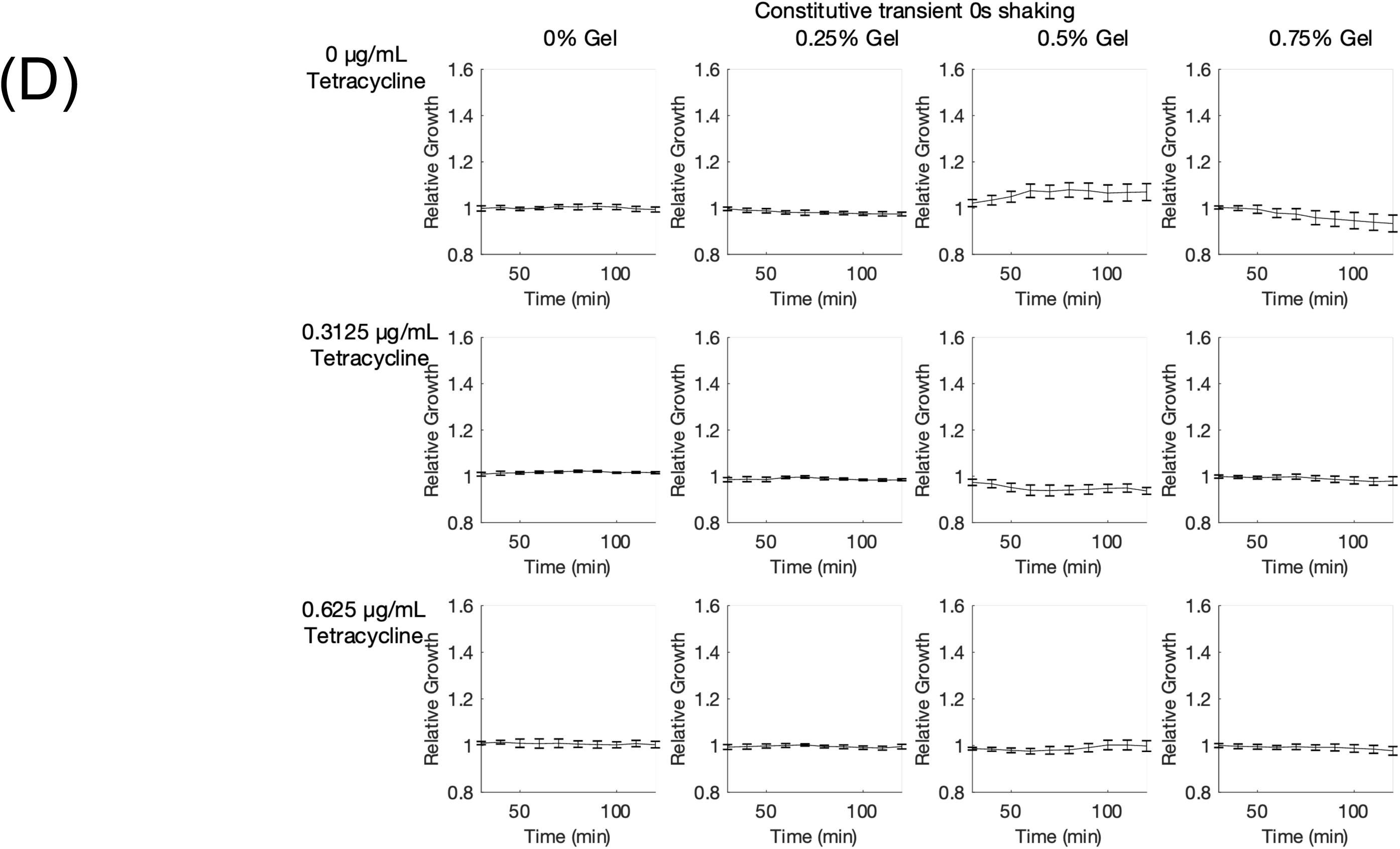
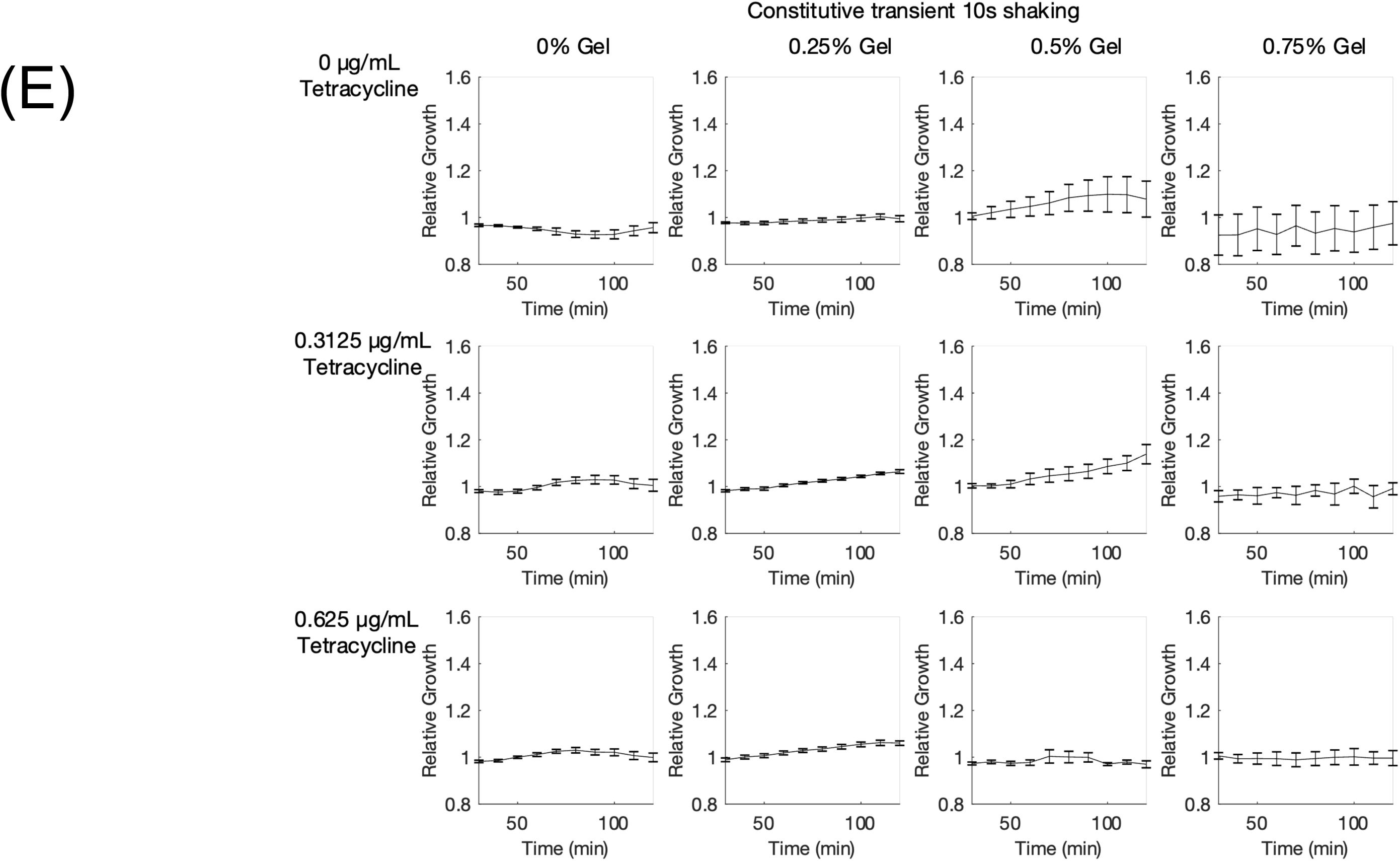
Changes in relative growth over time for DH5*α* cells expressing TetX grown in hydrogel and tetracycline. (A) Changes in relative growth over time for DH5*α* cells expressing TetX constitutively, grown without shaking. (B) Changes in relative growth over time for DH5*α* cells expressing TetX constitutively, grown with 10s/min shaking. Large reading at an early time point of 0.75% gel and 0 µg/mL tetracycline correspond to a bubble interfering with reading. (C) Changes in relative growth over time for DH5*α* cells expressing TetX constitutively, grown with 50s/min shaking. (D) Changes in relative growth over time for DH5*α* cells expressing TetX-ssra constitutively, grown without shaking. Degradation tag (ssra) makes TetX unstable. (E) Changes in relative growth over time for DH5*α* cells expressing TetX-ssra constitutively, grown with 10s/min shaking. Degradation tag (ssra) makes TetX unstable. (A-E) Cell grown in LB with 0, 0.25, 0.5, or 0.75% gel along with 0 µg/mL, 0.3125 µg/mL, or 0.625 µg/mL tetracycline. Points represent mean values, and error bars represent standard error of the mean. Mean and SEM are calculated using six biological replicates except for (C), which is calculated using four biological replicates.

**Supplementary Figure 8.**
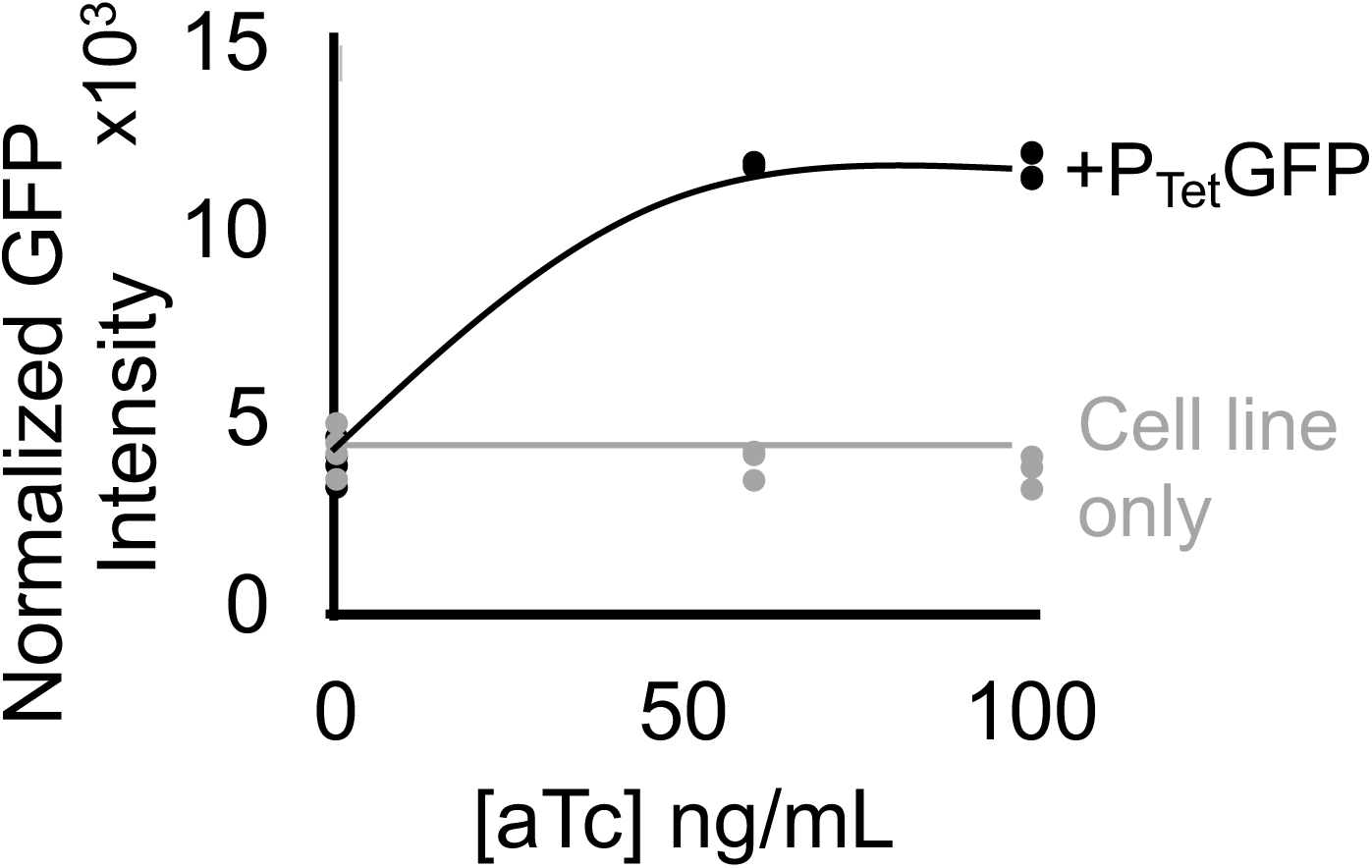
Inducible circuit characterization. Characterization of P_Tet_ producing GFP in DH5*α*pro cells in response to anhydrous tetracycline. Cell line only control is background fluorescence of DH5*α*pro cells. Points represent individual replicates, line highlights trend. There are four replicates per condition.

**Supplementary Figure 9.**
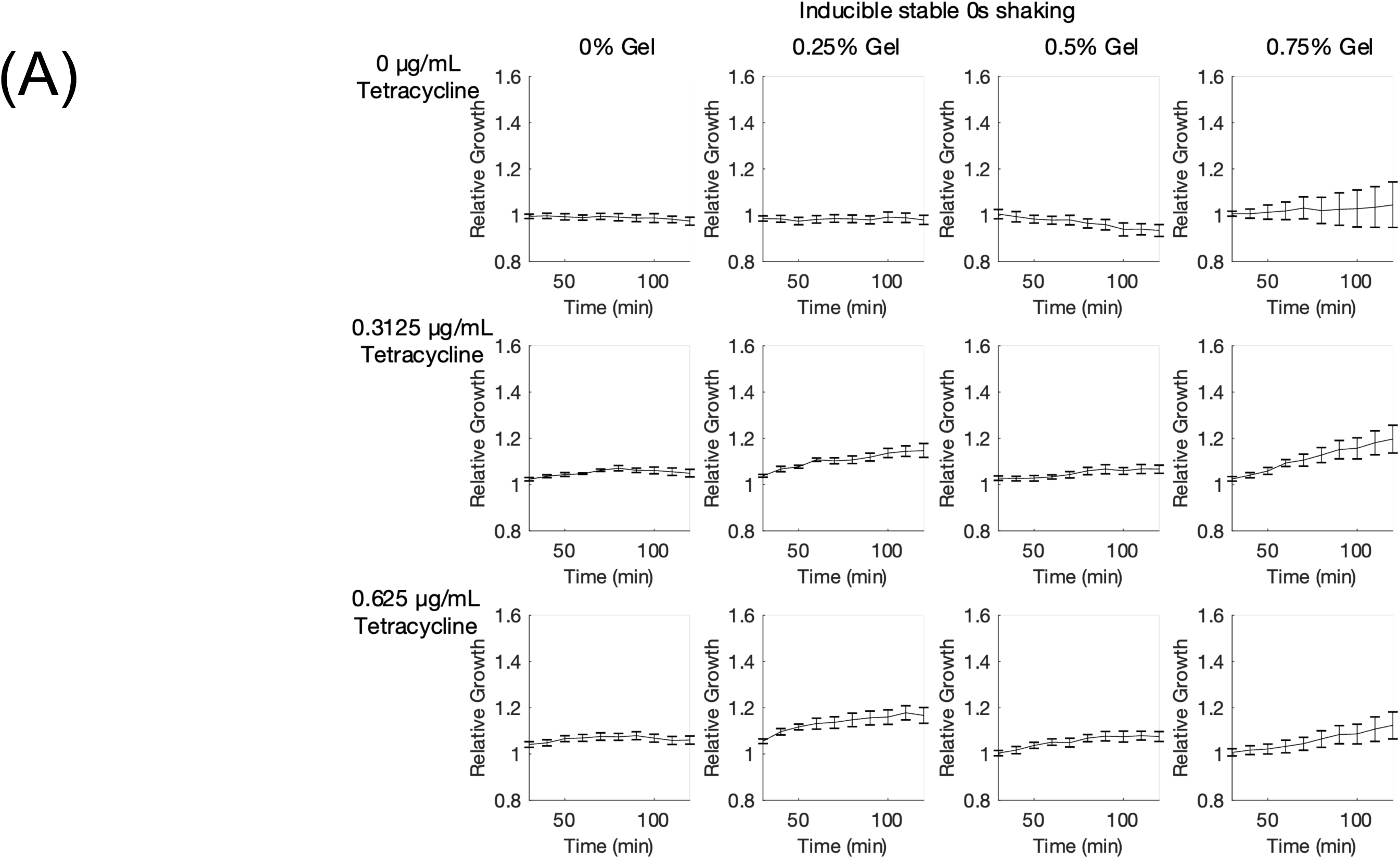
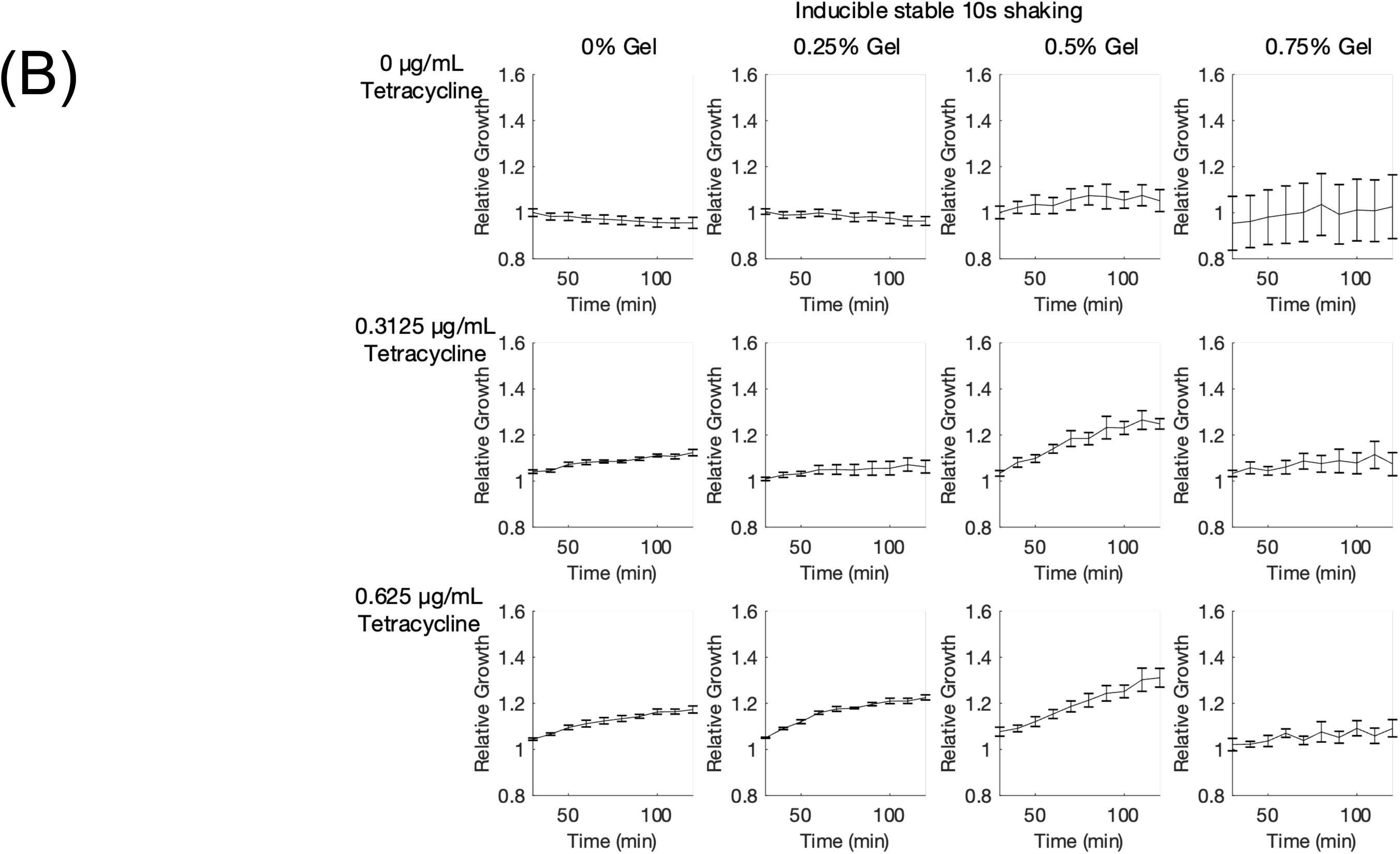
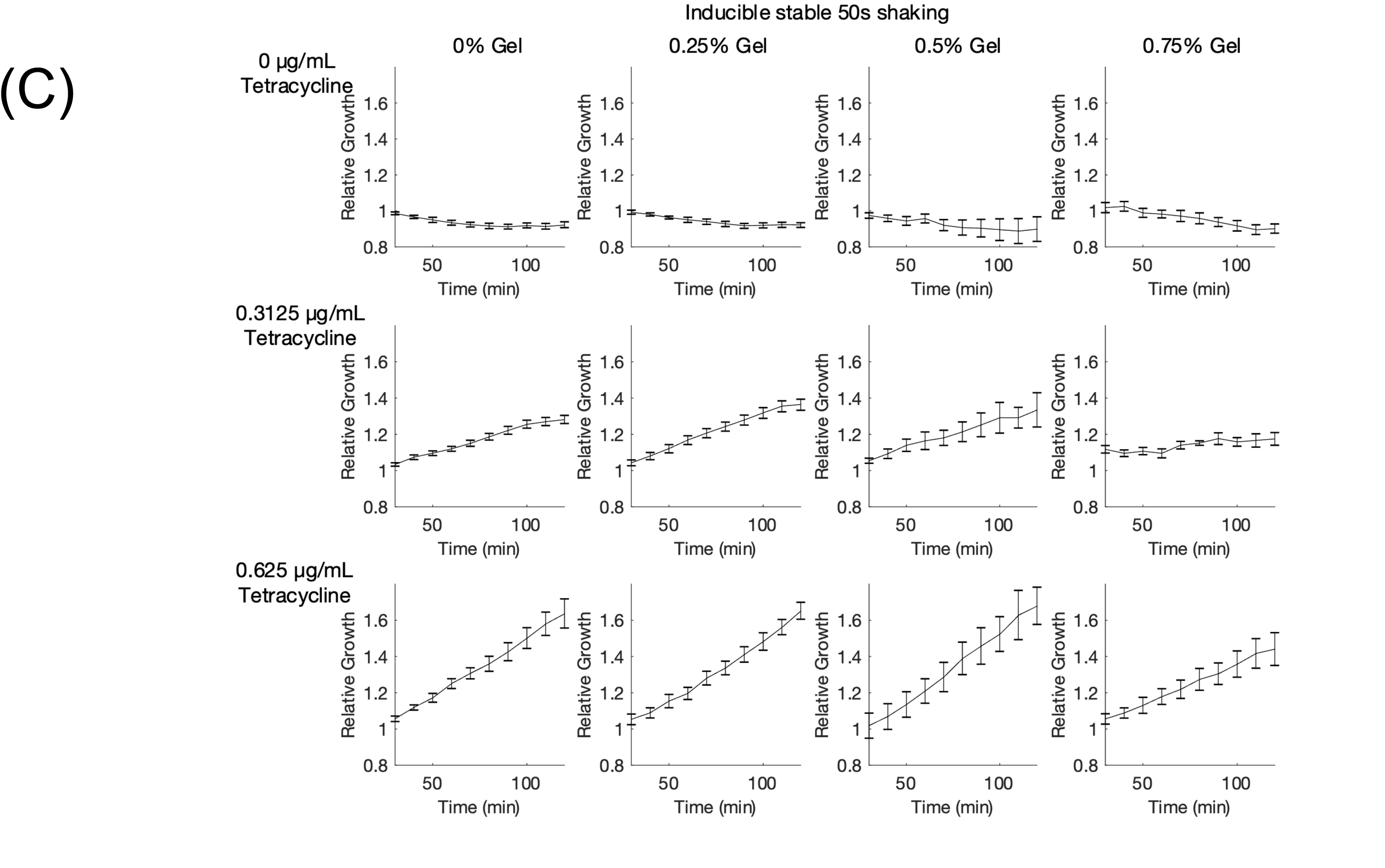
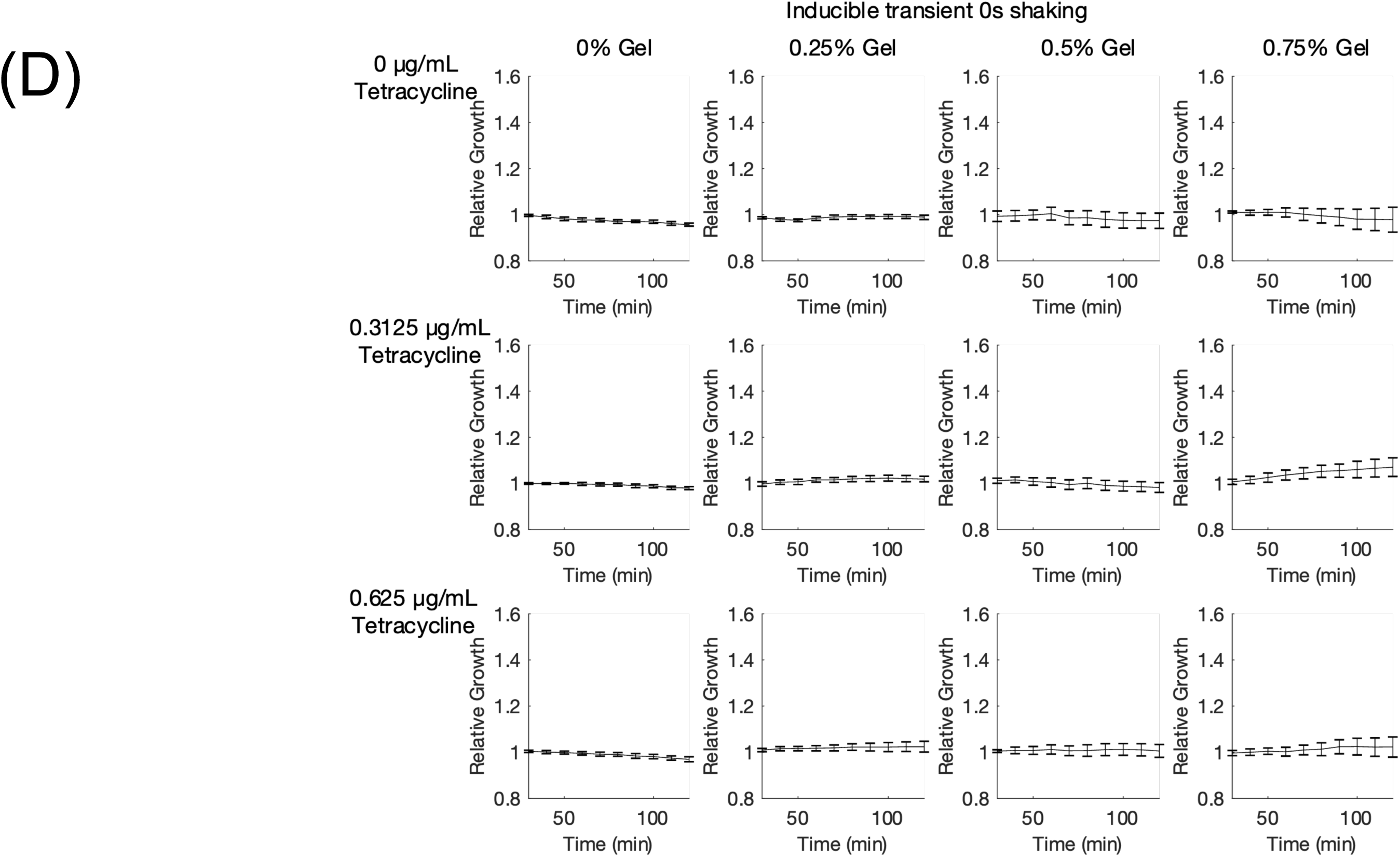
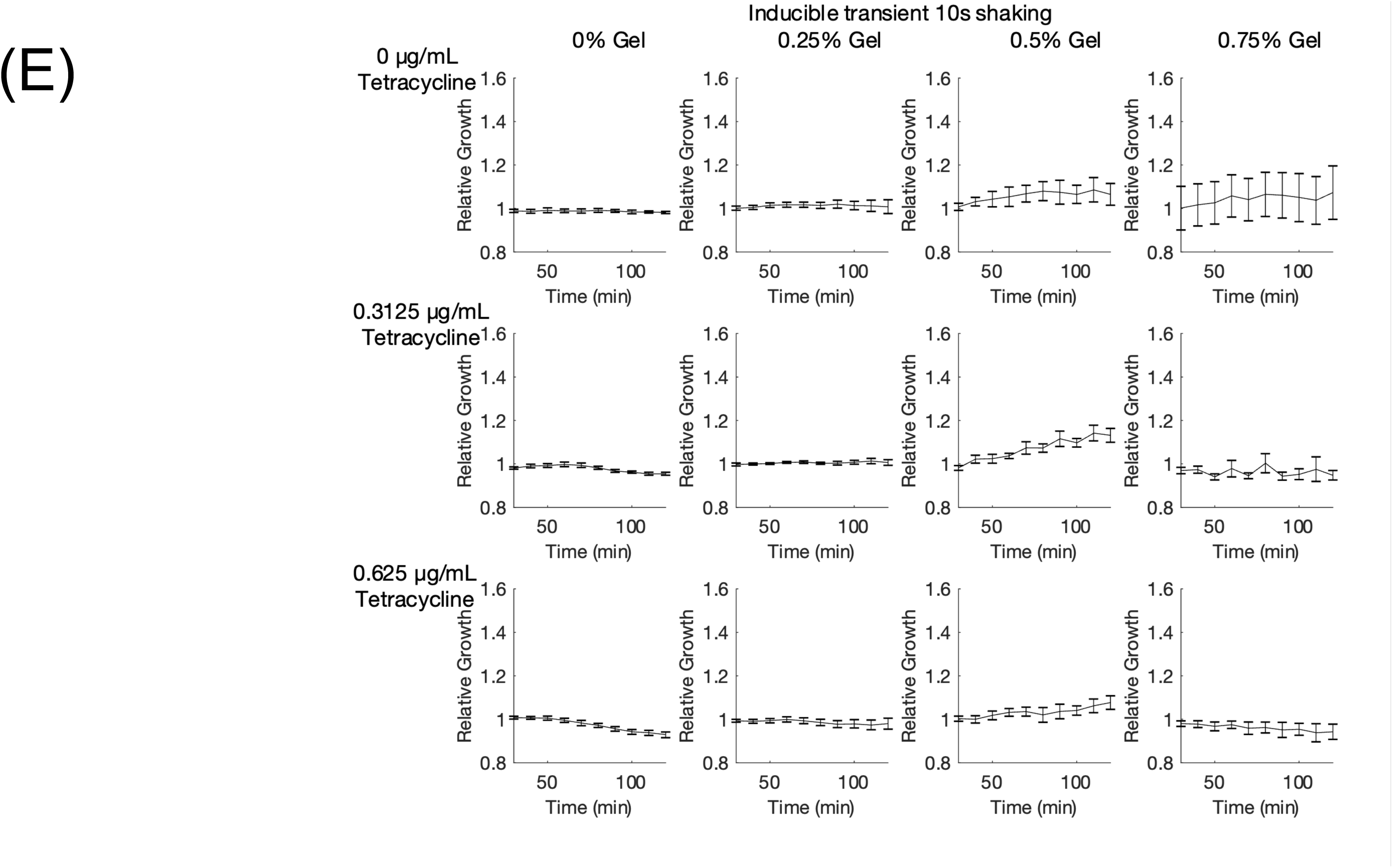
Changes in OD600 over time for DH5*α*pro cells expressing TetX grown in hydrogel and tetracycline. (A) Changes in relative growth over time for DH5*α*pro cells expressing TetX upon tetracycline induction, grown without shaking. (B) Changes in relative growth over time for DH5*α*pro cells expressing TetX upon tetracycline induction, grown with 10s/min shaking. (C) Changes in relative growth over time for DH5*α*pro cells expressing TetX constitutively, grown with 50s/min shaking. (D) Changes in relative growth over time for DH5*α*pro cells expressing TetX-ssra upon tetracycline induction, grown without shaking. Degradation tag (ssra) makes TetX unstable. (E) Changes in relative growth over time for DH5*α*pro cells expressing TetX-ssra upon tetracycline induction, grown with 10s/min shaking. Degradation tag (ssra) makes TetX unstable. (A-E) Cell grown in LB with 0, 0.25, 0.5, or 0.75% gel along with 0 µg/mL, 0.3125 µg/mL, or 0.625 µg/mL tetracycline. Points represent mean values, and error bars represent standard error of the mean. Mean and SEM are calculated using six biological replicates.

